# The Guinea Pig: A New Model for Human Preimplantation Development

**DOI:** 10.1101/2024.02.05.578945

**Authors:** Jesica Romina Canizo, Cheng Zhao, Sophie Petropoulos

**Author notes:** Corrresponding author.

## Abstract

Preimplantation development is an important window of human embryogenesis. During this time, the initial lineages are formed which largely govern embryo competence, implantation, and ultimately the developmental potential of the fetus. Ethical constraints and limitations surrounding human embryos research often necessitates the use of a model system. We now identify the guinea pig as a promising small animal model, which closely recapitulates early human embryogenesis in terms of the timing of compaction, early-, mid-, and late-blastocyst formation and implantation. We also observe conserved spatio-temporal expression of key lineage markers, roles of both Hippo and MEK-ERK signaling and an incomplete X-Chromosome inactivation. Further, our multi-species analysis highlights the spatio-temporal expression of conserved and divergent genes during preimplantation development. The guinea pig serves as an exciting new model which will enhance developmental and pluripotency research and can be leveraged to better understand the longer term impact of early exposures on offspring outcomes.

## INTRODUCTION

Human preimplantation development is a foundational period during which the initial three lineages are established, consisting of the trophectoderm (TE), the primitive endoderm (PE), and the epiblast (EPI)^1^. This window of preimplantation development is not only important for the establishment of these primary lineages, but provides the basis for subsequent embryonic development through the continued division and differentiation of cells during gastrulation and organogenesis. Further, perturbations to the molecular events and regulatory mechanisms, as observed with ‘insults’ or xenobiotics exposures, during this period might influence the developmental trajectories of various tissues and organs, shaping the phenotypic outcomes and long-term health of the individual. Furthermore, understanding the fundamental aspects of preimplantation development has implications for assisted reproductive technologies and infertility treatments, providing valuable insights for optimizing conditions that support healthy embryonic development. In essence, the study of preimplantation development is imperative for unravelling the intricate processes that lay the foundation for human life, encompassing both the immediate embryonic stages and their far-reaching consequences on overall health and well-being.

Due to the limited access and ethical restrictions surrounding research to work with human embryos, historically, understanding human embryology has heavily relied on mammalian model organisms^2^. While mouse studies have played a pivotal role in unveiling fundamental principles of early development^3^, recent advancements in low-input methods for investigating genetic and epigenetic mechanisms and efficient techniques for assessing gene function have led to the study of mammalian embryos across various species. This is vital for deciphering the molecular mechanisms involved in pluripotency and understanding how embryos adapt to maintain pluripotency through diverse developmental strategies and from an evolutionary perspective^4^. Such knowledge is essential for developing chemically defined culture media to preserve various pluripotent states in embryo-derived stem cell lines and to understand fundamental principles of early lineage development.

The guinea pig (*Cavia Porcellus*) has long been an established model for reproductive studies and generally shares remarkable similarities to humans in terms of general physiology^5^. In the context of reproduction and development, the guinea pig is the only laboratory rodent with a full estrus cycle, encompassing both follicular and luteal phases, similar to humans, cows, ewes, and pigs. This is in contrast to other laboratory rodents, such as mice, rats, and hamsters, which have incomplete and endocrinologically distinct estrus cycles lacking functional luteal phases^6^. In early development, the guinea pig preimplantation period is ∼6-7 days, similar to the human, representing one of the only animal models that parallels the duration of human preimplantation^7–10^. Similar to humans and in contrast to other animal models like mice, rats, rabbits, pigs, sheep and cows, the guinea pig undergoes interstitial implantation and cavitation^11^ and following implantation, both the guinea pig and human EPI undergo cavitation and form a bilaminar disc^12^. Post-implantation, the guinea pig placenta closely mirrors that of humans, as both species have haemomonochorial placentation and proliferating trophoblast cells of similar subtypes to those of the human^13,14^; making the guinea pig a good model for understanding trophoblast differentiation and amniogenesis in the human.. Finally, their gestation period of 68-72 days can be categorized into trimesters, mirroring the stages of fetal development in humans^15^ and the guinea pigs give birth to neuro-anatomically mature offspring^16^; representing an animal model that closely recapitulates early human development^10^.

Despite these notable similarities, the guinea pig preimplantation embryo has not been thoroughly characterized as the majority of studies have focused on the post-implantation period. Leveraging the guinea pig as an *in vivo* model to study fundamental mechanism(s) underlying these early embryogenesis milestones and how perturbations or exposures to certain drugs/xenobiotics/exogenous compounds impact these processes opens new avenues to enhance our understanding of these inaccessible events in the human. Recognizing the observed parallels in early development and general physiology between guinea pigs and humans, we aimed to characterize the guinea pig preimplantation embryo with the hopes of identifying both conserved and diverging aspects, ultimately introducing an additional small animal model which can be used for better understanding preimplantation development and pluripotency. We also wanted to identify a small animal model resembling human embryos that would enable us to examine longer-term phenotypic consequences of preimplantation exposures/’insults’ on fetal development and offspring outcomes.

To address this gap, we conducted a comprehensive characterization of preimplantation development in the guinea pig. We determined the morphokinetics and characterized the temporal development of lineage formation using immunofluorescence in parallel with single-cell RNA sequencing, identifying key genes and transcription factors that govern blastocyst formation. In addition, we assessed the role of aPKC and the evolutionarily conserved Hippo signalling pathway on blastocyst formation and MEK-ERK signaling on PE formation. Moreover, using H3K27me3 loci as a proxy for X-Chromosome activity and RNA FISH for long noncoding RNA *XIST*, we compared the status of the X-Chrs in female mouse, human, and guinea pig embryos. Finally, we used a comparative biology approach to highlight similarities and differences in early development amongst the guinea pig, mouse, and human with a focus on lineage development and naive pluripotency, highlighting the importance of cross-species studies Overall, our study identifies the guinea pig as a promising new small animal model which can be leveraged to better understand the molecular underpinnings that govern human preimplantation development and by extension, gastrulation.

## RESULTS

### Characterization of Preimplantation Development in the Guinea Pig

Very little is known about the guinea pig preimplantation embryo. We aimed to characterize and stage the embryo based on morphokinetics to determine the timing of key events during this window and establish the spatiotemporal expression of key lineage markers for trophectoderm (TE), inner cell mass (ICM), epiblast (EPI), and primitive endoderm (PE). Embryos were collected following the presence of a positive sperm smear starting at 82-84 hrs post-fertilization, or embryonic day (E) 3.5 corresponding to the 8-cell stage, continuously at 3-6 hr intervals until we were unable to flush in vivo embryos (approximately E6). The inability to flush embryos past E6 corroborates previous work demonstrating that implantation occurs between E6-E7 in the guinea pigs^17^, and is similar to that observed in humans.

We first took bright-field images of each embryo collected and then utilized a combined approach based on embryo morphology, cell number, and embryonic day for embryo staging (Fig. 1a). Compaction was identified through visual inspection of cell boundaries, occurring between the 8 and 16 cell stage, consistent with prior reports in humans^18^. We then leveraged human embryo time-lapse data acquired by Meistermann et al.^19^, and similarly aligned compaction (E4-E4.5, 16 cell stage (C)) with T = 0 h. Notably, the morphokinetics observed in the guinea embryos exhibited temporal progression akin to that seen in humans. The transition from compacted morula to precavitation occurred at T = 9 h, followed by cavitation, early blastocyst (EB) occurring at T = 21 h, mid-blastocyst (MB) at T = 27 h and late blastocyst (LB) at T = 33 h post-compaction (T = 31 h in the human)^19^ (Fig. 1a).

**Figure 1.**
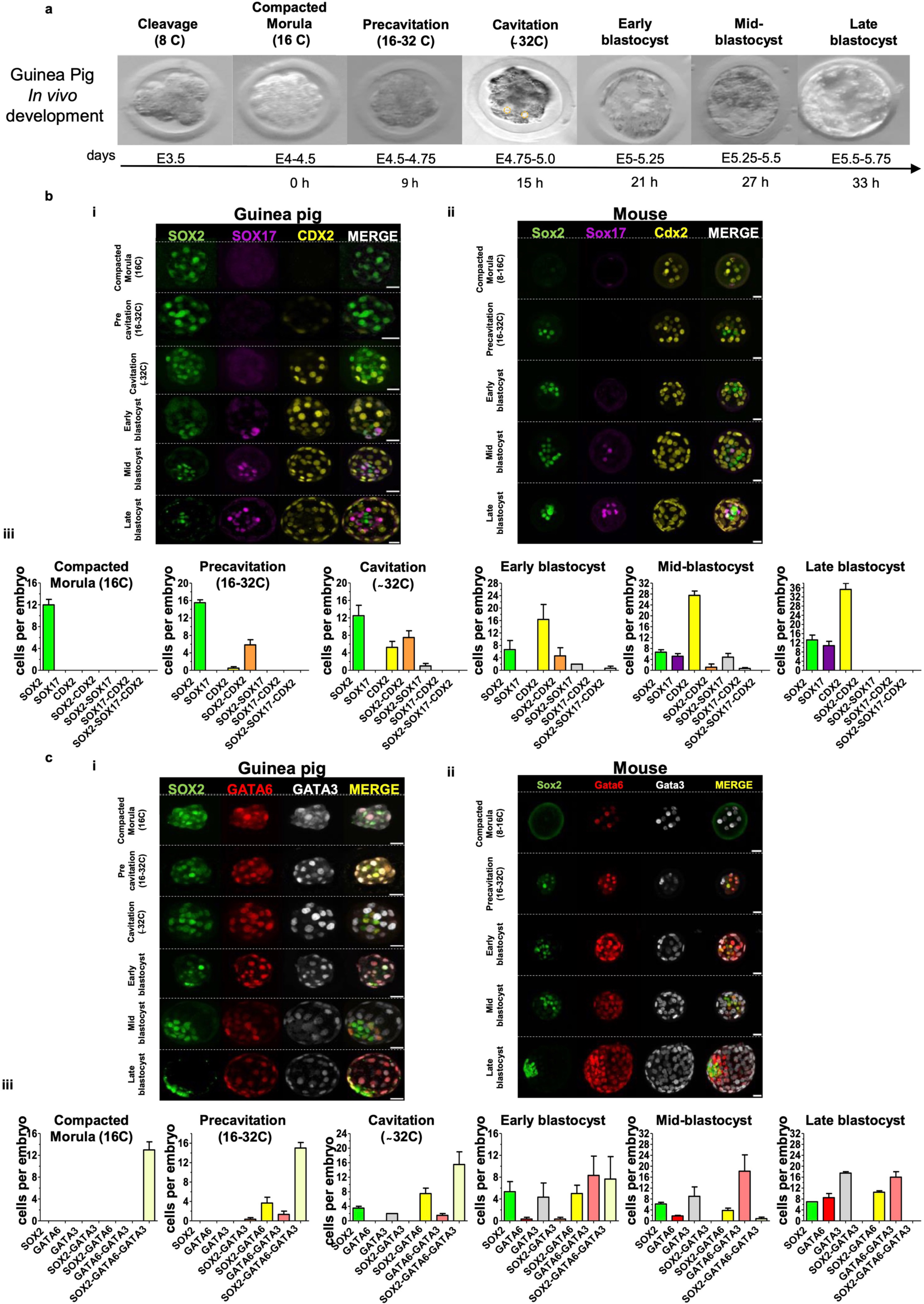
Time-course characterization of *in vivo* guinea pig preimplantation development. (a) Bright-field images of embryos captured *in vivo* at different time points, highlighting the link between developmental stages, morphological phases, embryonic days (E), and the average number of cells observed. At the cavitation stage, microlumens are marked by yellow dotted lines. (b) Time-course immunofluorescence analysis of EPI (SOX2), PE (SOX17) and TE (CDX2) associated markers in guinea pigs (i) and mouse (ii) embryos; bar graphs (iii) showing the quantification of the number of cells per guinea pig embryo of SOX2, SOX17 and CDX2 only positive cells as well as their co-expression at different developmental stages: compacted morula (n = 3), pre-cavitation (n = 6) and cavitation (n = 4), early (n = 3) mid (n = 6) and late blastocysts (n = 4). (c) Immunofluorescence analysis of EPI (SOX2), PE (GATA6) and TE (GATA3) associated markers in guinea pig (i) and mouse (ii) embryos; bar graphs (iii) showing the quantification of the number of cells per guinea pig embryo of SOX2, GATA6 and GATA3 only positive cells as well as their co-expression at different developmental stages: compacted morula (n = 4), pre-cavitation (n = 3) and cavitation (n = 2), early (n = 3) mid (n = 6) and late blastocysts (n = 3). Data are presented as Mean ± SEM Scale bars: 20 µm.

Next, we wanted to determine the spatio-temporal expression of key molecular markers for the lineages. Using a comparative biology approach, we performed parallel experiments in mouse embryos and leveraged previously published data from human embryos^20,21^. Starting at the compacted morula stage (E4-E4.5, 16C), we determined that the guinea pig embryo expressed ubiquitous expression of SOX2 and lacked SOX17 and CDX2 (Fig. 1bi), similar to the human embryo at this stage of development^20^. In contrast, in the mouse, Sox2 expression was restricted to the inner cell population with Cdx2 exclusively localized to outer cells (Fig. 1bii), consistent with previous studies^22,23^. Moreover, at this stage, all cells in guinea pig embryos expressed SOX2/GATA6/GATA3 (Fig. 1ci), aligning with previous reports in humans^21^. In contrast, in the mouse, distinct inner and outer cells were marked with Sox2 or Sox2+Gata6 and Gata3, respectively (Fig. 1cii). Collectively, the expression patterns of these markers indicate that lineage specification does not commence in the compacted guinea pig embryo, aligning with observations in human, rat, and bovine^3^.

CDX2 is a downstream effector of the Hippo signalling pathway, and in the mouse is expressed starting at the late 8-cell embryo and is a key gene driving TE formation (Fig. 1bii). In contrast, CDX2 emerges in the human, bovine and rat embryo around the time of cavitation^3,21^. Here we see the progressive embryo maturation and distinct expression of CDX2+ emerging in outer cells during the 16C-32C stage, which corresponds to E4.5-E4.75 (Fig. 1bi and biii and Extended Data Fig. 1ai-ii). Recognizing the dynamic nature of embryo development, we considered not only developmental time/embryonic day but also cell number and the presence and size of a blastocoel compartment for staging, consequently categorizing embryos within the 16C-32C stage as precavitation and cavitation. During precavitation, we observed a dynamic range of CDX2+ expression (3-10 cells) in outer cells with the majority of the embryos coexpressing SOX2+/CDX2+ suggesting that the TE program is being poised during this time but not initiated (Fig. 1bi and biii and Extended Data Fig. 1a). In two embryos collected during this stage, distinct CDX2+ cells were observed in only 1-2 of the outer cells (Extended Data Fig. 1 ai and aii), followed by a drastic increase in exclusive expression of CDX2+ and a simultaneous decrease of SOX2+ in the outer cells at cavitation, suggesting initiation of the first lineage segregation, ICM-TE, in the guinea pig. The timing of this initial lineage specification is reinforced by the emergence of cells only expressing GATA3+ during cavitation E4.75-5 in guinea pig embryos (Fig. 1ciii). In humans, CDX2 is upregulated in E5 blastocysts and initially coincident with OCT4, indicating a lag in CDX2 expression in the TE lineage ^20^, relative to the mouse and as we observe in the guinea pig. Moreover, during cavitation, we captured two embryos (n = 2/4) expressing 2 cells of SOX2+SOX17+ indicating that the PE program may begin to be poised after the emergence of ICM-TE (Extended Data Fig. 1bi). In guinea pigs, the transition from precavitation to cavitation is rapid, approximately 6 h, underscoring the importance of sequential sampling during this developmental phase to capture ongoing events. This rapid transition is similar to the leading model of lineage specification in humans, where around E5, embryos shift from an absence of defined lineage to an intermediate ICM-TE and a subsequent emergence of EPI/PE, all occurring during the same developmental day.

During the early blastocyst stage (EB, E5-E5.25, 28-36 cells with the presence of an early blastocoel), the number of cells co-expressing SOX2 and CDX2 is reduced, as we observe the continued ramping up of CDX2+ outer cells, representing a more defined TE lineage. We also begin to see an increase in the expression of SOX17 and a subpopulation of cells solely expressing GATA6 (Fig. 1bi, 1biii, 1ci and 1ciii). These findings suggest the emergence of the second lineage segregation, consisting of the transition from ICM to EPI/PE. By mid-blastocyst (E5.25-E5.5, 40-50 cells with a more enlarged blastocoel), the specification of the three lineages is more pronounced indicated by the increased number of cells solely expressing one of the well-established molecular marks for their corresponding lineage and the decreased number of cells co-expressing these markers (Fig. 1bii, 1biii, 1cii, 1ciii). In the mouse during this window (Fig. 1bii), Sox17 is turned on and coexpressed with Sox2 in a salt and pepper pattern in the ICM, marking a contrast with what is observed in the guinea pig and human, where SOX17 expression is initiated earlier^20^. By the late blastocyst stage (LB, E5.5-E5.75, 55-66 cells with an expanded large blastocoel), just prior to implantation, lineage specification is complete in guinea pigs and there are no longer the presence of cells co-expressing lineage markers. This aligns with previous reports in humans that demonstrate well-defined lineage segregation before implantation^19,20,24^.

### Characterizing the Preimplantation Guinea Pig Embryo using Single-cell RNA sequencing

A limitation of immunofluorescence is the number of lineage markers which can be simultaneously assessed. To obtain a comprehensive overview of the spatial-temporal gene expression and lineage dynamics throughout guinea pig preimplantation development, we utilized single-cell RNA sequencing (scRNA-seq) on individual blastomeres isolated from E3.5-E6 embryos. After quality control (see Supplemental Methods), we retained 541 high-quality single-cell transcriptomes from 42 embryos, with 19311 genes with at least 1 read in one cell, a minimum of 3,054 genes expressed per cell, and 87.1 % mapping ratio (Fig. 2a and Extended Data Fig. 2, Supplementary Table 1).

**Figure 2.**
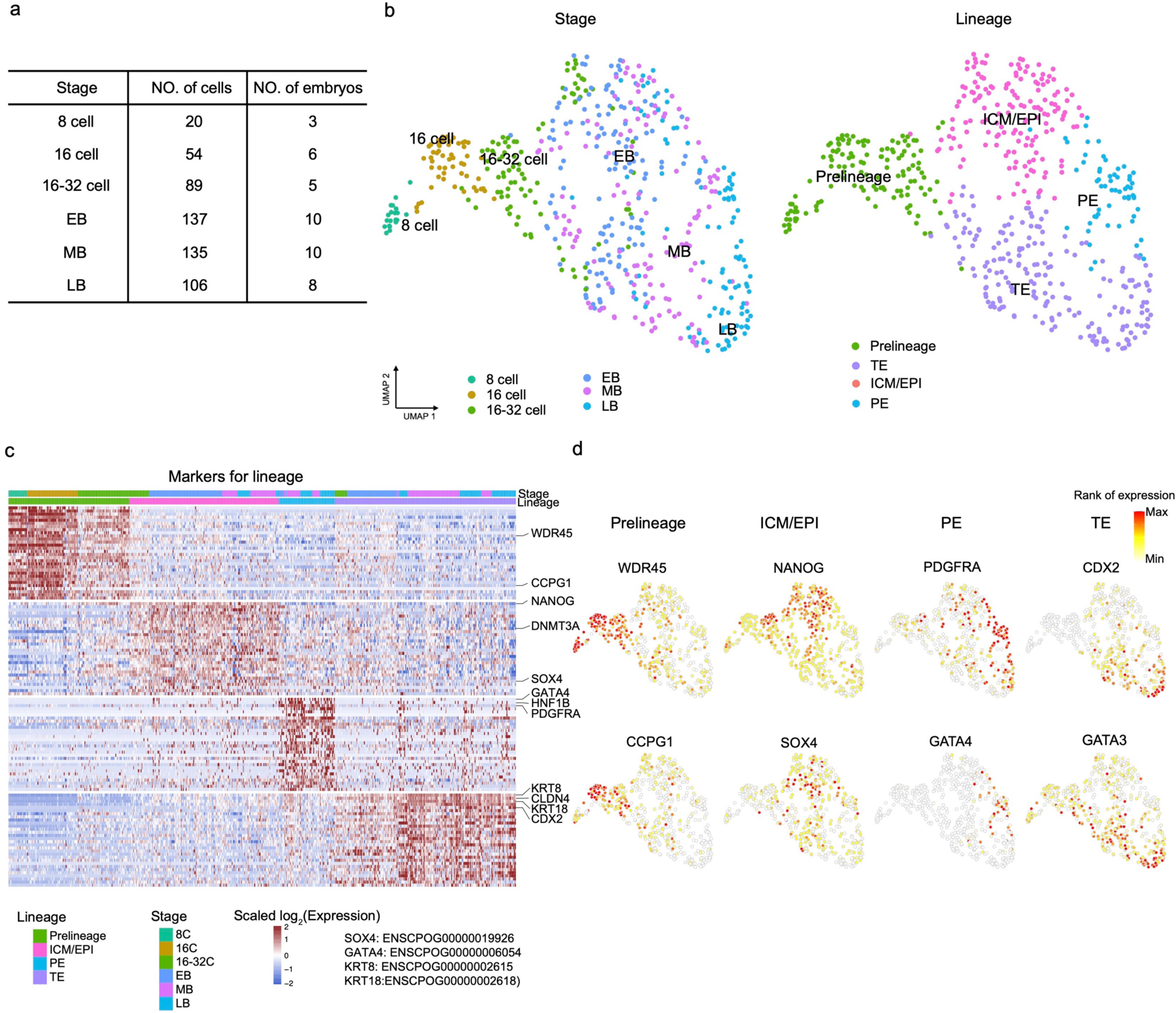
Characterization of preimplantation development in the guinea pig with single-cell RNA-sequencing. (a) Number of cells belonging to each lineage included in the analysis. (b) Two-dimensional UMAP representations of 541 single-cell transcriptomes from guinea pig preimplantation embryos. Lineages and developmental stages were indicated by colors, respectively. (EB: early blastocyst, MB: middle blastocyst, LB: late blastocyst). (c) Heatmap showing the top 30 marker genes for Prelineage, EPI, PE, and TE. Selected well-known lineage genes labeled on the right. (d) UMAP plot of cells showing selected marker gene expression. Prelineage genes include *WDR45* and *CCPG1*, ICM/EPI genes include *NANOG* and *SOX4*, PE genes include *PDGFRA* and *GATA4* and TE genes include *CDX2* and *GATA3*.

To better characterize the preimplantation embryo and assign cell identity, based on gene expression, we performed dimensionality reduction using Uniform Manifold Approximation and Projection (UMAP), providing a clear visualization of embryo development. Development time accounts for the primary factor driving expression variation and includes 8 cell (8C), 16 cell (16C), and 16-32 cell (16-32C; which includes both precavitation and cavitating embryos), EB, MB, and LB (Fig. 2b and Extended Data Fig. 3a-c). Next, we identified the lineage populations present by leveraging well-known markers from human and mouse studies for each lineage (TE, ICM, EPI, and PE)^19,24–27^ and assigned cell identity as prelineage (encompassing the 8C, 16C and precavitation 16-32C cells), TE, ICM/EPI, and PE (Fig. 2b-d and Extended Data Fig. 3c). Notably, within this dataset and consistent with human studies^19,24,26^, we were unable to distinguish a unique cluster of cells corresponding to the ICM (from resolution 0.4 to 1.4 using “FindCluster” function, Extended Data Fig. 3d) and as such have labeled the lineage as ‘ICM/EPI’.

To confirm the identity of the cell populations and obtain gene signatures unique to each guinea pig lineage, we performed marker detection for the six populations (Supplementary Table 2). Lineage identification was verified by observing the expression of known candidate markers in the human embryo, such as WD Repeat-Containing Protein 45 (*WDR45*) and Cell Cycle Progression Protein 1 (*CCPG1*) for prelineage (8C to precavitation inclusive), Nanog Homeobox (*NANOG*), Sry-box Transcription Factor 4 (*SOX4*) for ICM/EPI, platelet-derived growth factor receptor alpha (*PDGFRA*), GATA Binding Protein 4 (*GATA4*) for PE, and Caudal Type Homeobox 2 (*CDX2*), GATA Binding Protein 3 (*GATA3*) for TE (Fig. 2c, d).

### Timing of Lineage Specification in the Guinea Pig and ICM Resolution

To confirm the timing of the first lineage specification (ICM-TE), we determined the developmental trajectory of all cells using the top 450 DEGs obtained and a two-dimensional diffusion map with consisting of 8C, 16C, and the precavitation 16-32C, which we call ‘prelineage’ as they occur before the bifurcation point of the ICM/EPI and TE cells (Fig. 3a, b). Next, maintaining the assigned cell pseudotimes, we checked the distribution of cells stratified by embryo stage and observed that the ICM-TE segregation occurs during E4.5-5, corresponding to the transition from precavitation to cavitation (Fig. 3c) and in alignment with our immunostaining findings from Fig. 1. Examining the top 10 DEG per lineage branch we observed an upregulation of a well-known genes related to TE as *CDX2*, and a dramatic reduction in *NANOG* expression in the later TE lineage branches and simultaneous increase of *NANOG* in the later ICM branches (Fig. 3d). Notably, *IL17RD*, a gene encoding for an interleukin receptor is also upregulated in the ICM branch. This receptor negatively regulates FGFR signaling by limiting MAPK signaling via its intracellular domain in xenopus and zebrafish embryos^28,29^. *DNMT3B*, a *de novo* DNA methyltransferase, is highly expressed in both ICM/EPI and TE lineages, suggesting the potential for global hypermethylation during embryo development, similar to what is observed in human embryos^30^.

**Figure 3.**
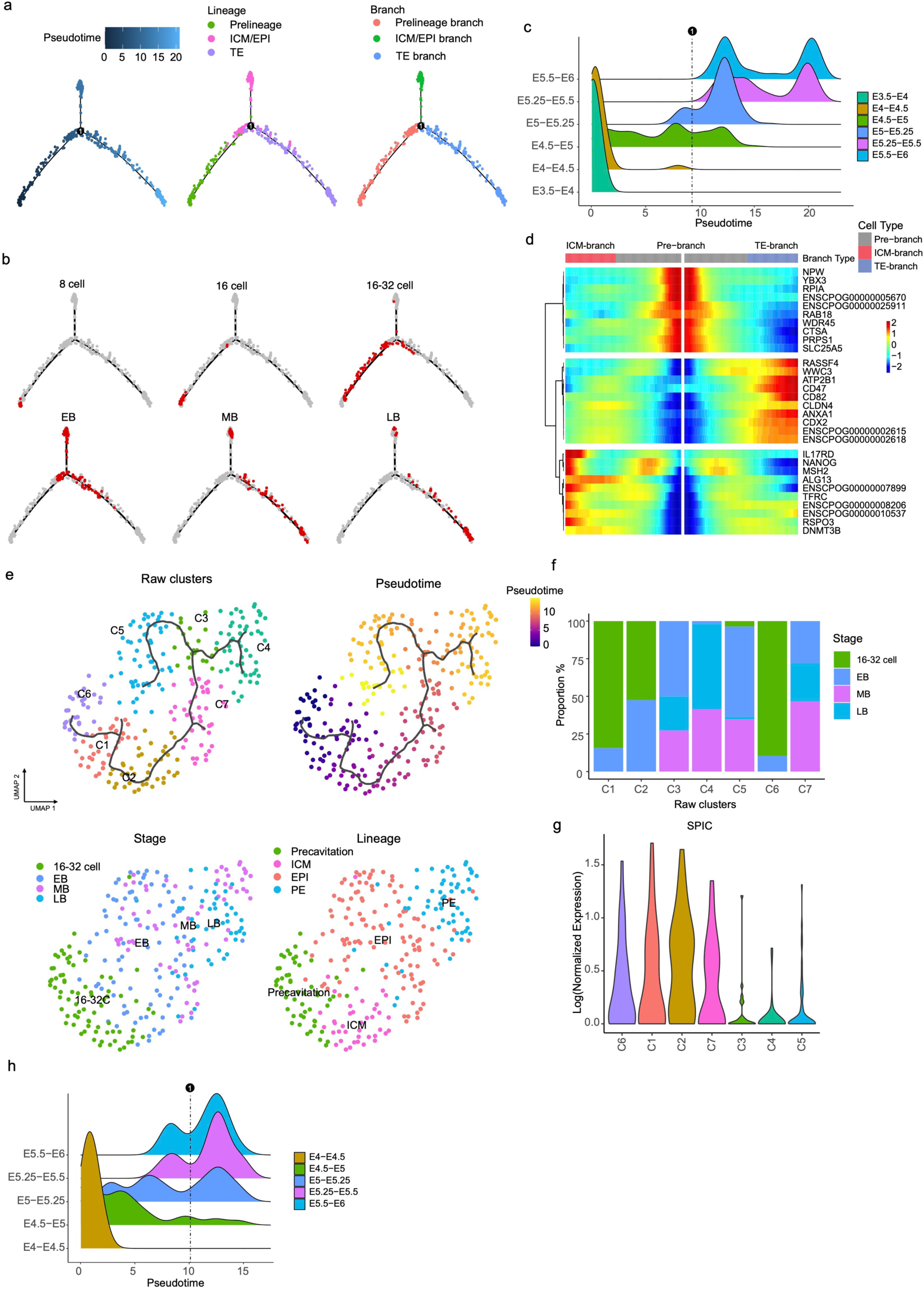
Timing of ICM/TE and EPI/PE segregation. (a) Two-dimensional diffusion map showing the developmental trajectory of prelineage, ICM/EPI, and TE cells. Pseudo-time and lineages were indicated by colors, respectively. (b) Highlighted cells from different developmental stages on the trajectory shown in (a). (c) Ridge plot showing the distribution of single cells along the pseudo time axis stratified by embryonic days. The dashed line on the ridge plot represents the bifurcation into TE and ICMs. (d) Heatmap representing the top 10 DEGs among Prelineages, ICM/EPI, and TE.. (e) Two-dimensional UMAP representations cells from precavitation, ICM, EPI, and PE cells. Raw clusters, Pseudo-time and stages and lineages were indicated by the colors, respectively. (f) Bar plot representing the proportion of stages in each cluster from (e). (g) Violin plots illustrating the expression of SPIC in each cluster from (e). (h) Ridge plot showing the distribution of single cells along the pseudo-time axis, stratified by embryonic days. The dashed line on the ridge plot represents the bifurcation into EPI and PEs.

We next interrogated the ICM/EPI population to determine whether we could identify a gene signature that was distinct to a subpopulation of these cells, representing the ICM. To resolve this, we subset the cells belonging to the precavitation, ICM/EPI, and PE populations, and performed clustering and pseudo-time inference using the top 1500 variable genes amongst them (Fig. 3e). Seven clusters were identified (Fig. 3e). Careful examination of the distribution of stages and inferred pseudotime, revealed that the majority of cells in clusters 3, 4, 5, and 7 belonged to the MB and LB stages (Fig. 3e, f). Based on immunostaining, we knew that “ICM” cells do not persist into the MB stages (cluster 7), where we already observed a defined EPI and PE. Therefore, we identified cluster 2 to represent the "ICM" since 52.5 % of the cells belonged to the “16-32 cell” and 47.5 % of the cells belonged to “EB” but was located before the bifurcation point of the EPI and PE branches. To validate this, we utilized known ICM specific markers recently identified in the human embryo including *SPIC*^31^. *SPIC*, is a marker of ground state pluripotency and conserved transcription factor in the ICM^31,32^. In the mouse and human, *SPIC* is highly expressed in the prelineage and ICM and then drastically declines in the EPI based on our reanalysis of published datasets^24,33,34^. In the guinea pig we observed high expression of *SPIC* in clusters 6, 1, 2, and 7, with a sharp drop in expression in clusters 3, 4, and 5. This drop in expression coincided with the bifurcation point during which the specification of EPI and PE occurred, confirming the identification of the "ICM" as cluster 2 (Fig. 3g). After checking the distribution of cells inferred by pseudotime, stratified by embryo stage, EPI and PE specification occurred during the E5-E5.25 corresponding to the EB (Fig. 3h). These analyses corroborated our immunostaining that progressive segregation of the ICM into EPI and PE occurs during the EB.

Finally, we analyzed the expression patterns of genes related to human naive and primed pluripotency states, as well as the well-known core markers of pluripotency in ICM, EPI, PE, and TE, to chart pluripotency following blastocyst formation. We observed a signature of naive and core pluripotency in guinea pig embryos similar but not identical to what is seen in human embryos (Extended Data Fig. 4). Furthermore, throughout blastocyst development, genes tied to primed pluripotency were very lowly expressed or absent in both species. With similar expression patterns for *DNMT3B* and *PRDM14* in both species, implying a conserved role in the embryo. The overlapping pluripotency signatures between guinea pig and human embryos underscore potentially conserved mechanisms related to the establishment and progression of the different states of pluripotency. Further exploration in this direction could provide valuable insights into the broader understanding of pluripotency across diverse species.

### Stage and Lineage-Specific Signalling Pathway Enrichment

To attain biological insight into the functionality of the cell populations, we determined the lineage-specific gene expression modules associated with each cell population, using Self-organizing maps (SOMs)^35,36^ followed by the pathway analysis using gene sets from the guinea pig hallmark collection in the Kyoto Encyclopedia of Genes and Genomes (KEGG) database^37^ (Fig. 4a and b, Supplementary Table 3). Early in development, at the 8-cell stage, we see an enrichment of known pathways involved in embryonic stem cell maintenance and preimplantation embryo development including “Wnt signalling pathway” (P = 0.016), “Signalling pathways regulating pluripotency of stem cells” (P = 0.021), “MAPK signalling pathway” (P = 0.012) and “Hippo signalling pathway” (P = 0.012). Just prior to cavitation, metabolism-related pathways such as “Tyrosine metabolism” (P = 0.030), and “Histidine metabolism” (P = 0.030) are enriched, potentially coinciding with the change in metabolic needs of the embryo as it transitions blastocyst formation (Fig. 4b, Supplementary Table 3). In the TE lineage, genes related to “Galactose metabolism” (P = 0.025) and “Glycine, serine and threonine metabolism” (P = 0.0053) as well as adhesion-related pathways such as “Gap junction” (P = 0.028), Focal adhesion (P = 0.00070) and “Cell adhesion molecules” (P = 0.00037) were enriched, supporting the important role of the TE for implantation. In addition, we observed an enrichment of `PI3K−Akt signalling pathway in both PE and TE lineages` (P = 0.027 and 0.0037 respectively) (Fig. 4b and Supplementary Table 3).

**Figure 4.**
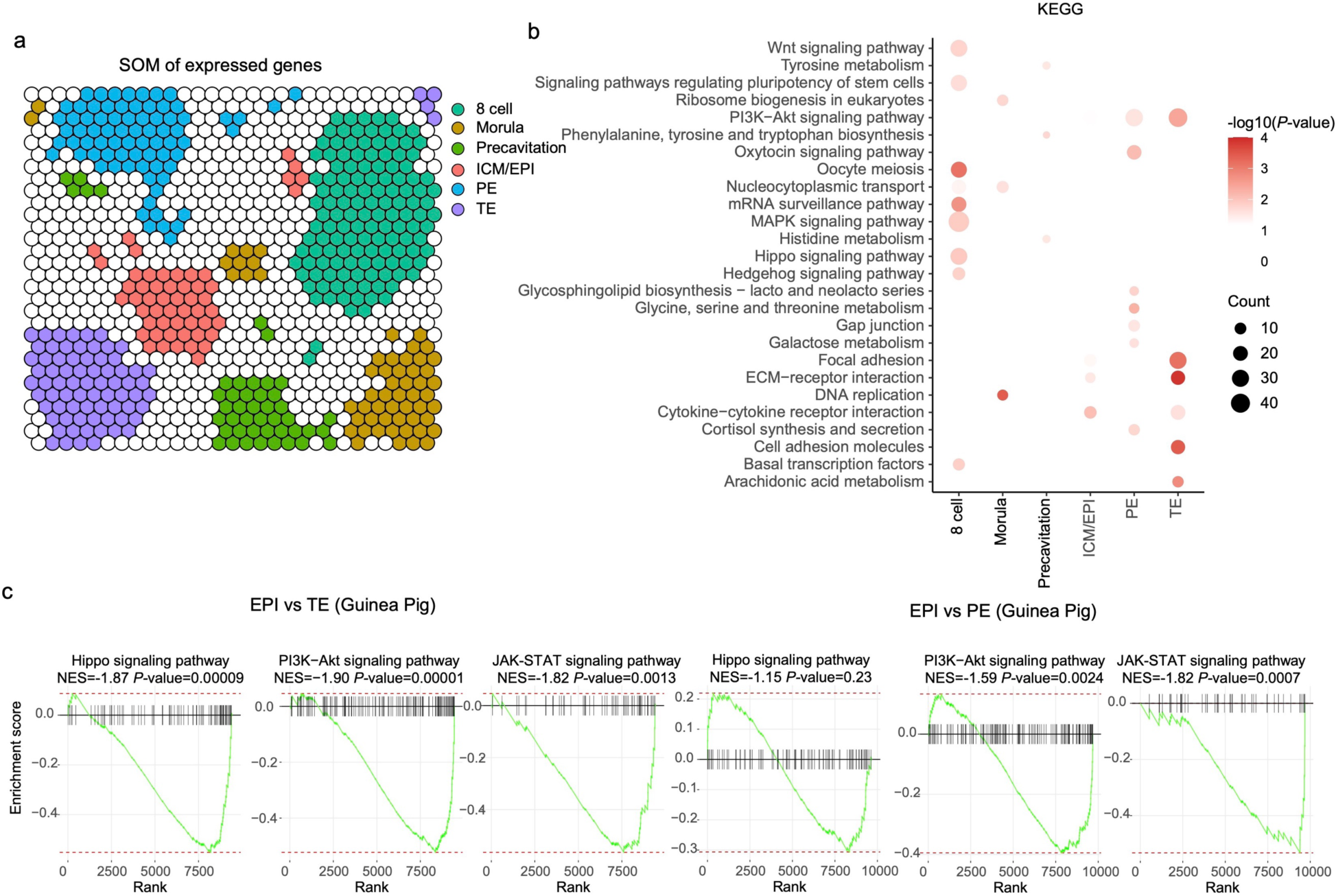
Signalling pathways related to preimplantation embryo development and lineage segregation. (a) Self-organizing map (SOM) of lineages based on guinea pig scRNA-seq expression, with lineage-specific clusters (Z-score > 1.5) represented by colors. (b) Dot plot presenting enriched KEGG pathways of lineage-specific gene modules, with size and color denoting the number of genes and significance (p-value) of the pathway. (c) Plots showing enrichment scores for genes with at least a 0.1 log2(Fold change) difference related to the "PI3K-Akt signalling pathway," "Hippo signalling pathway," and "JAK-STAT signalling pathway" in guinea pig EPI vs TE and EPI vs PE comparisons. A higher enrichment score indicates over-representation of the related gene set, while a negative score indicates under-representation.

Next we wanted to further explore the potential functional roles of signalling pathways during embryo development. Simultaneously considering lineage and developmental stages, the contribution of each pathway to gene expression variability was assessed through gene set co-regulation analysis (GESECA) (Extended Data Fig. 5). An upregulation of genes linked to DNA replication at the 8-cell stage was observed. Studies in mouse embryos have hinted at a connection between DNA replication and the onset of transcription during ZGA^38^. The disruption of nucleosomes during DNA replication could create an opportunity for maternally derived transcription machinery to access previously inaccessible promoters, potentially enabling the reprogramming of gene expression patterns and mark promoters for transcription during ZGA. Further, at the 8-cell, 16-cell, and precavitation stages, guinea pig embryos showed an overrepresentation of genes associated with the “Pentose phosphate pathway” (Extended Data Fig. 5a and b). Our findings suggest that the enrichment of the pentose phosphate pathway, which is crucial for nucleotide biosynthesis, coupled with the simultaneous upregulation of DNA replication genes around the 8-cell stage, implies embryonic genome activation in guinea pig embryos, akin to human embryos at this stage^2^; though we did not directly examine this in the guinea pig. Moreover, using the GESECA analysis, genes related to cysteine and methionine metabolism were overexpressed in EPI and TE lineages, but expressed at lower levels in pre-lineages, suggesting a role blastocyst formation (Extended Data Fig. 5). Previous studies in bovine^39^ and mouse^40,41^ preimplantation embryos demonstrated the significance of disrupted methionine metabolism, leading to impaired blastocyst formation. Our results suggest a potentially conserved role of methionine in the morula to blastocyst transition. We also observed an enrichment of "ECM-Receptor Interaction" and "Focal Adhesion" in both PE and TE lineages, possibly related to extraembryonic maturation before implantation (Extended Data Fig. 5a and b), in addition to pathways like "Rap1 Signaling Pathway" and "Insulin Signaling Pathway" (Extended Data Fig. 5a and b). Functional studies in guinea pig embryos will provide insights into the role of these pathways in blastocyst development.

Next, we checked pathways which may be involved in lineage segregation and/or maintenance, focusing on comparisons between EPI vs TE and EPI vs PE (Fig. 4c). We found Hippo signalling pathway genes were notably highly expressed in TE (normalized enrichment score (NES) = −1.87, P = 0.00009) compared to EPI, supporting previous findings that highlight the evolutionarily conserved role of Hippo signalling in TE formation^3^. Additionally, genes related to "JAK-STAT Signaling Pathway" and "PI3K-Akt Signaling Pathway" are overall highly expressed in TE compared to EPI (P-values of 0.00001 and 0.0013, respectively) and in PE compared with EPI (P-values of 0.0024 and 0.0007). This suggests that the activation of JAK-STAT and PI3K-Akt signalling is essential for pluripotency loss and the formation of PE and TE lineages in guinea pig embryos. There is currently a lack of studies in the human embryo examining the JAK-STAT pathway, making a direct comparison not possible, but this aligns with findings in porcine embryos, where JAK-STAT signalling activation by IL6 has a critical role during pig TE specification^42^. Of note, the predicted function of PI3K-AKT signalling in guinea pigs is in contrast to that observed in porcine^42^ and human^43^ embryos, where PI3K-AKT activation via IGF1 promotes ICM proliferation by increasing pluripotent cells. Investigating whether PI3K-AKT function differs between guinea pigs and human embryos requires further exploration and may provide additional mechanistic insights into blastocyst formation, lineage segregation, and potential evolutionarily conserved or divergent roles.

### Determining the Presence of Mural and Polar TE Sub-Lineages

In the human embryo, the polar TE initiates implantation into the uterine wall, while in the mouse and guinea pig, it is the mural TE that initiates attachment and implantation^19,44^. Despite the presence of mural-polar TE sub-lineages in guinea pig embryos, their molecular signatures remain unexplored. As the TE plays a unique role in mammalian development by forming the embryonic portion of the placenta ^45^, we aimed to investigate the presence of mural-polar sub-lineages and their corresponding gene signatures. First selected all TE cells from cavitation to LB stages and performed UMAP dimensional reduction, leading to the identification of two distinct clusters (Fig. 5a). Following differential expression analysis between the two clusters (Fig. 5b), ‘TE sub-lineage 1’ displayed 4 up-regulated genes overlapping with the human mural TE, while “TE sub-lineage 2” displayed 21 genes overlapping with the human polar TE. For each TE sub-lineage, two genes (*ERBB3* and *NUCB2* in sub_lineage 1 and *PTGES* and *SLBP* in sub_lineage 2) were identified which displayed the opposite enrichment between the two species (Fig. 5b). To validate the localization of each TE sub-lineage in the guinea pig, we then selected the human polar marker, Nuclear Receptor subfamily 2 group F member 2 (NR2F2). In the guinea pig, NR2F2 was localized to the mural TE sub-lineage (Fig. 5b). Notably, in the human, the polar sub-lineage constitutes the smaller proportion of TE cells adjacent to the ICM. Thus, we aimed to determine the relative contribution of the TE sub-lineages in the guinea pigs by quantifying the number of NR2F2+/GATA3+ (mural) and GATA3+ only cells (polar) (Extended Data Fig. 6a). In contrast to the human, we observed that the dominant population of cells in the TE corresponds to the polar lineage.

**Figure 5.**
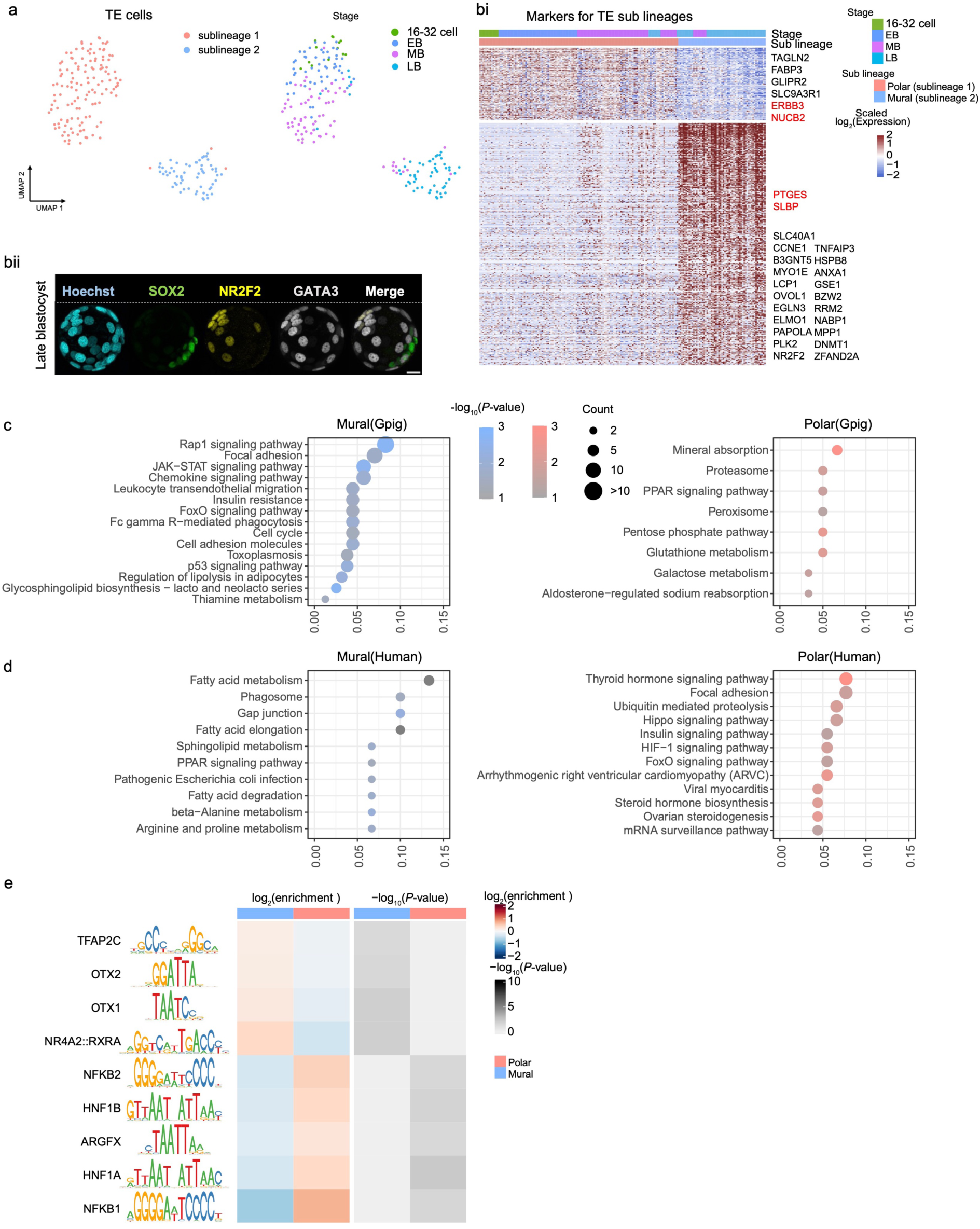
Profiling the Mural and Polar TE Sub-lineages. (a) Two-dimensional UMAP displaying the sub-lineages in guinea pig TE cells. Sub-lineages and developmental stages are indicated by colors, respectively. (bi). Immunofluorescence image of a representative *in vivo* guinea pig embryo at the late blastocyst stage labeled with Hoechst (nucleus), SOX2+ (EPI), NR2F2+GATA3+ (mural TE) and GATA3+ (polar TE). (bii) Heatmap representation displaying the differentially expressed genes between TE sub-lineage cells. Genes that were also significantly differentially expressed between human mural and polar TE cells are labeled on the right. Red and black colors indicate the same and opposite expression patterns. (c) Dot plot illustrating the enriched KEGG pathways for DEGs between mural and polar TE cells in guinea pig. Color and size indicate the significance and number of differentially expressed genes in each pathway. (d) Dot plot illustrating the enriched KEGG pathways for DEGs between mural and polar TE cells from Human. Color and size indicate the significance and number of differentially expressed genes in each pathway. (e) Identification of significantly enriched motifs located in the promoter region of DEGs between guinea pig mural and polar cells. The enrichment and P-values of listed motifs were displayed in the heatmap.

To explore functionality associated with each TE sub-lineage, we conducted KEGG pathway enrichment analysis on the significant DEGs between mural and polar cells and observed an upregulation of the “PPAR signalling pathway” (P = 0.022) and metabolism-related pathways, including Glutathione and Galactose metabolism (P = 0.0068 and P = 0.023, respectively) in polar cells (Fig. 5c). Additionally, an increase in “Chemokine signalling pathways” (P = 0.012) and “Cell adhesion molecules” (P = 0.012) was noted in the mural, aligning with their role in implantation (Fig. 5c, Supplementary Table 4). Considering the substantial overlap between guinea pig mural-enriched genes and human polar-enriched genes, we sought to identify enriched pathways in human polar TE. Conducting a similar KEGG analysis on DEGs between human mural and polar cells revealed the enrichment of the PPAR signaling pathway and FoxO signaling pathway, mirroring the findings in guinea pig mural cells (Fig. 5d).

Next, we wanted to further examine the TE sublineage signatures between the human and guinea pig. To do this, we included any mural-polar DEGs in either the human or guinea pig giving us 615 genes in total (Extended Data Fig. 6b). We observed that 360 of these genes were enriched in the opposite lineage between guinea pig and human (Extended Data Fig. 6b). Functional enrichment analysis underscored the conservation of genes related to "Glycosphingolipid biosynthesis" (P = 0.021), "Tight junction" (P = 0.0074), and "Adherens junction" (P = 0.022), crucial for embryo implantation into the uterine wall (Extended Data Fig. 6c). Intriguingly, genes associated with "Fatty acid metabolism" (P = 0.0037), "Insulin resistance" (P = 0.0025), "Focal adhesion" (P = 0.00019), and "Glutathione metabolism" (P = 0.00049) exhibited opposite expression patterns between human and guinea pig, hinting at potential metabolic differences between the mural and polar cells in these species.

Finally, we performed an enrichment analysis of transcription factor motifs within promoters of DEGs between guinea pig mural and polar cells (Fig. 5e). Although an FDR less than 0.05 was not achieved, likely due to the lower number of DEGs, we did identify biologically relevant motifs, such as TFAP2C, part of the core regulatory circuitry of the TE program^46^ and NR4A2:RXR, nuclear receptors regulated by retinoids and steroids and thus may be important for priming the TE for implantation^47^ (Fig. 5e).

### Cross-Species Comparison

To better assess both similarities and differences between guinea pig and human embryos, we employed the anchor identified by Canonical Correlation Analysis (CCA) to identify conserved cell types amongst the datasets^48^. By integrating our guinea pig transcriptome with publicly available human embryo single-cell RNA-seq datasets^24,49^, we visualized the combined data in a 2-dimensional UMAP space (Fig. 6a). The cells representing prelineage, ICM, EPI, PE, and TE aligned consistently across both species (Fig. 6a). Subsequently, a Mutual Nearest Neighbor (MNN) search was conducted to establish equivalence in developmental time between the cells of human and guinea pig preimplantation embryos (Fig. 6b and c). Guinea pig cells aligned with their counterparts in humans for the cell lineage and developmental stage, supporting our immunostaining findings (Fig 6b and c). Generally, a closer relationship is observed amongst cells spanning the 8-cell stage to precavitation (Fig. 6b).

**Figure 6.**
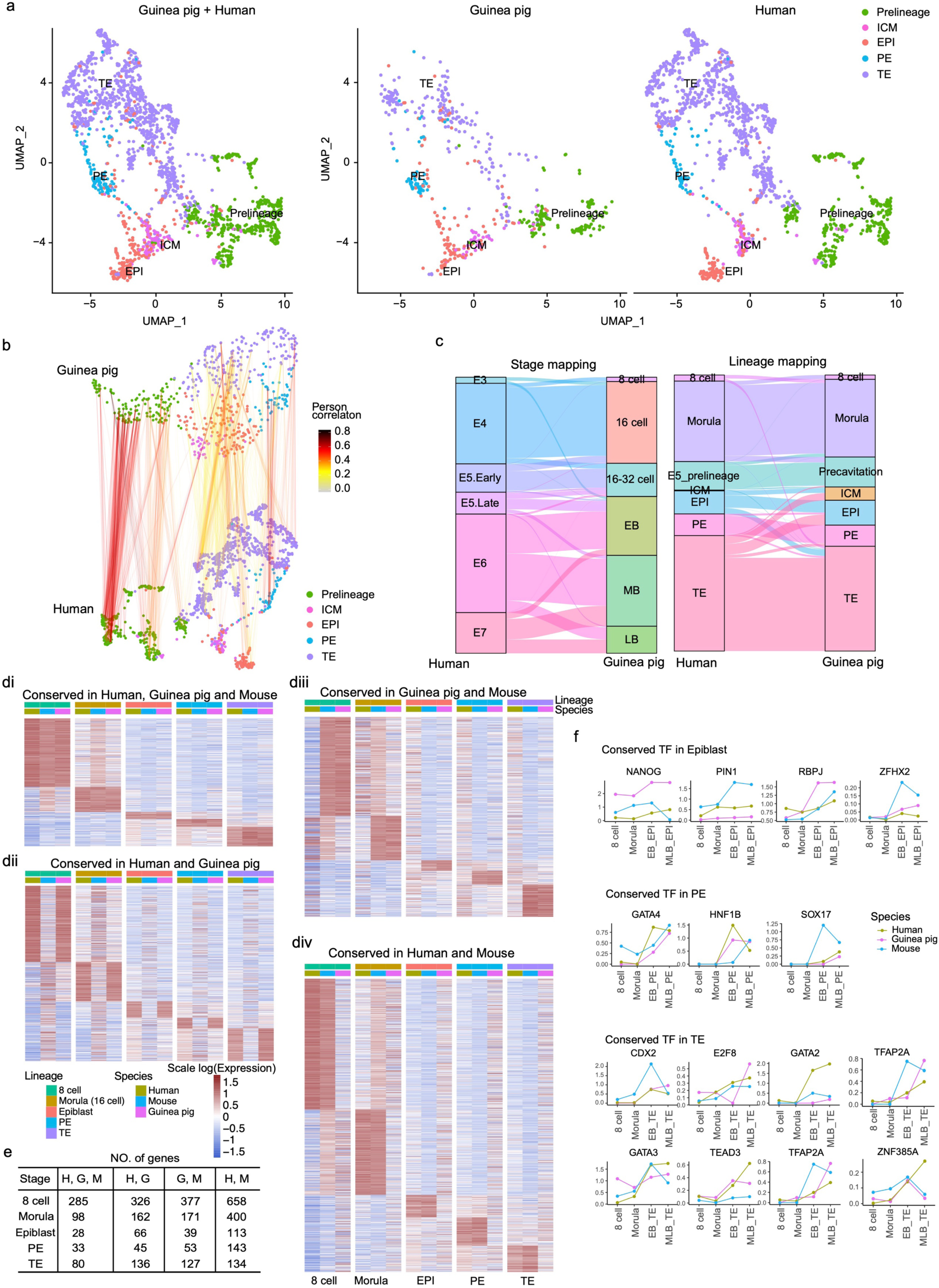
Cross-species analysis. (a) Integration of preimplantation single-cell RNA data from human and guinea pig. (b) Subspace projection illustrating the linkage between mutual nearest neighbor pairs in human and guinea pig. Line colors indicate Pearson correlation coefficients. (c) Alluvial plot comparing guinea pig developmental stages and lineages with human embryonic days and lineages among mutual nearest neighbour pairs. (d) Heatmap representing conserved genes within a lineage amongst species: (i) human, guinea pig, and mouse; (ii) human and guinea pig but not in mouse; (iii) guinea pig and mouse but not in human and (iv) human and mouse but not in guinea pig. (e) Number of genes categorized in different groups in (d). (f) Line plot displaying the expression of selected transcription factors stratified by stages and lineages in human, guinea pig, and mouse. Different species are represented by different colors.

Next, we performed a multi-species analysis to identify similarities and differences amongst the mouse, human, and guinea pig preimplantation embryos (Fig. 6d and e). To enrich our cross-species analysis, we incorporated two additional preimplantation mouse datasets^33,50^. Given that expression of genes in embryonic cells are not always on/off amongst the lineages, we restricted our analysis to consider a gene within a specific lineage if it was the highest expressed in that particular lineage when compared to the others (see Methods). We first identified conserved genes for each lineage amongst the different species combinations (Supplementary Table 5). Overall, 524 genes were found to be conserved amongst the mouse, human and guinea pig lineages (Fig. 6di). These included, in EPI cells genes encoding for *FGF4*, *NANOG*, *MAP4K1*, *TDGF1* and *USP28*; in PE cells genes encoding for *GATA4*, *SOX17*, *HNF1B*, *DENND2C* and *PDGFRA* and in TE cells genes encoding for *LRRFIP1*, *KRT18*, *GATA3*, *GATA2*, *KRT19*, *KRT8* and *CLDN4* (Supplementary Table 5). From the 524 genes, we then selected the transcription factors to determine their expression pattern to consider both lineage and developmental time (Fig. 6f). We observed interesting cross-species patterns for *NANOG*/*Nanog* (peak expression observed in all three species at mid- to late-blastocyst), *SOX17*/*Sox17* (peak expression in the guinea pig and human at mid- to late-blastocyst), *CDX2*/*Cdx2* (no expression in the guinea pig and human at the 16-cell stage), and *TFAP2A*/*Tfap2a* (peak expression observed for both the guinea pig and human at mid- to late-blastocyst). These discrepancies in expression patterns highlight the importance of not only considering lineage, but also developmental time when examining regulatory factors in preimplantation development (Fig. 6f).

We next wanted to examine genes that are conserved in only two of the three species in different combinations. In total, 745 genes were conserved between the human and guinea pig but with discrepancy (either absent or expressed in a different lineage) in the mouse (Fig. 6dii and Supplementary Table 5). These included *SEPTIN6* and *LRP4* in EPI cells; *BMP2*, *TRIM, FGFR1* and *FGFR2* in PE cells; *DLX3*, *CD24*, *DAB2*, *EZR*, *NR2F2* and *PPARG* in TE cells. Next we examined genes with conserved peak expression between the guinea pig and mouse but not in the human, suggesting a rodent specific signature (Fig. 6iii and Supplementary Table 5). In total 767 genes were identified including *UHRF1, USP9X,* and *ACP6* in EPI; *F2R, ARF5, ERP29* and *FGF10* in PE cells; and*, ELF5, WNK2, KRT7* and *EOMES* in TE cells. Notably, *ELF5* and *EOMES*, previously identified as TE genes exclusively enriched in mouse TE and absent in human TE^24,51^, are also enriched in guinea pig TE suggesting a conserved role in rodent TE. Finally we identified 1,448 genes with conserved peak expression between the mouse and human lineages but not the corresponding lineage in the guinea pig (Fig. 6d iv and Supplementary Table 5). These include among others, in EPI cells genes encoding for *ETV1, UTF1, FBP1* and *GDF3*; in PE cells genes encoding for *FOXA2, COL4A1* and *IGF1*; and in TE cells genes encoding for *S100A6, TACSTD2, TDG, TPM4* and *ADK*.

Finally, from a comparative biology approach to obtain a more comprehensive multi-species comparison, we plotted the expression of selected well known lineage markers for the human, marmoset, cynomolgus, mouse, and guinea pig (Extended Data Fig. 7). Generally, primates, non-human primates and rodents demonstrated a remarkable conversation in the expression of these genes but with some striking differences in timing. For example, at the morula stage, zinc finger and SCAN domain containing 4 (*ZSCAN4*), involved in telomere maintenance and embryonic genome activation (EGA), was detected only in the human embryos in our analysis (Extended Data Fig. 7). However, in mice there are nine paralogous genes and one orthologous gene in the marmoset reported^52^. Previous reports have noted exclusive expression of *Zscan4* at the 2-cell stage mouse embryo^52^, consistent with mouse EGA. This highlights differences in temporal gene expression related to the timing of developmental milestones and again emphasizes the importance of taking into account both lineage and time when performing cross-species analyses. Furthermore, while the expression of the PAIRED (PRD)-like homeobox gene Arginine-Fifty Homeobox (*ARGFX*) and the double homeobox A (*DUXA*) appears to be exclusive to primates, the DUX-family seems to have a conserved role in placental mammals^53^. It is important to note that the absence of a gene in the guinea pig may be attributed to incomplete genome assembly rather than biological absence. In mouse, *Duxf* is expressed at thee early 2C stage and is implicated in EGA with redundant function alongside *Obox4*^54^. In primates other than humans, genes from the DUX family have also been identified^55^, but their expression in embryos and functional role in EGA remain to be elucidated. Consequently, there is a possibility that we might be underestimating the transcriptional similarities between guinea pigs and other species. *SPIC*, a marker of ground state pluripotency and conserved transcription factor in the ICM^31,56^ was found conserved between primates and rodents. Of note, the cynomolgus dataset lacks the 8-cell and morula timepoints, where *SPIC* may be highly expressed. In all primates and rodent blastocysts, bone morphogenetic protein 2 (*BMP2*) expression peaks in the PE, however, *PDGFRA* and *GATA4* are conserved across species. In the EPI cells, peak *FGF4* expression is conserved amongst all species. In contrast, peak expression of *TGFB3* is only observed in the human EPI. TGFβ signaling plays a role in expanding the pluripotent inner cell mass in humans embryos, as evidenced by the decrease in NANOG and SOX17 when TGFβ is inhibited^25,51^). This is in contrast to marmoset monkey embryos, where no effect is observed after treatment^25^. In mice, where Nodal is expressed later in epiblast maturation, TGFβ inhibition has no effect on Nodal, Sox17, or Oct4 expression at late blastocyst stages^51^ suggesting a difference in timing and function. In guinea pigs, the expression pattern of *TGFβ3* suggests a role in PE and TE, though this remains to be explored. Finally, for *TFAP2C,* peak expression is observed during both the 8-cell stage as well as the TE across all species. Consistent with this, *TFAP2C* is a distinctive feature of human naive stem cells and preimplantation EPI cells^25,26,57^ as well as a well known master regulator of TE formation^46,58,59^.

### Functional Analysis of Signaling Pathways

An evolutionary conserved molecular cascade initiates the TE program in human, cow, rat and mouse embryos at the morula stage^3,21^, when outer cells undergo apico-basal cell polarity acquisition, marked by the enrichment of the polarity protein complex at the contact-free domain. Atypical protein kinase C (aPKC) forms part of this complex to establish cell polarity and functions upstream of the Hippo signaling pathway^3,21^. Given the conserved role of both Hippo signaling and aPKC, we wanted to assess the impact of PKC inhibition on guinea pig and mouse embryos by administering the Gö6983 inhibitor starting from the compacted morula until the blastocyst stage using a dose determined from our dose-response experiments (Extended Data Fig. 8ai-iii and 9ai-ii). When both guinea pig and mouse embryos were allowed to progress to the blastocyst stage, control embryos formed expanded blastocysts, while embryos treated with the PKC inhibitor were arrested at cavitation (Fig. 7ai-ii and Extended Data Fig. 9).

Next, we investigated the role of Hippo signaling kinases (LATS1/2) in the guinea pig compared to mouse embryos using the specific LATS inhibitor, TRULI^3,60^. A dose-response experiment (see Methods for ranges) was conducted to determine the optimal dose of TRULI for guinea pig (Extended Data Fig. 8bi-iii)and mouse embryos (Extended Data Fig 9bi-iii). We determined that 5 µM and 7.5 µM of TRULI was the optimal dose for the mouse and guinea pig respectively. Mouse optimal dose is the same as a previously published work^3^. Both compacted guinea pig (16-cell) and mouse (8-cell) embryos were cultured with either TRULI (7.5µM or 5µM, respectively) or DMSO-control until the blastocyst stage (Fig. 7b and Extended Data Fig. 10a). In mouse embryos, consistent with previous reports^3,61,62^, Lats1/2 inhibition led to a complete ablation of Sox2 expression (in 5/6 embryos) and ectopic expression of Gata3 and active Yap (aYap) in the inner cells (Extended Data Fig. 10a). In contrast, in the guinea pig we observed a trend toward downregulation of SOX2 accompanied by ectopic expression of GATA3 and aYAP and (Fig. 7b-iii, P < 0.001), mirroring reports in human embryos^3^. Additionally, in guinea pig blastocysts, the number of outer cells expressing aYAP and GATA3 remained unaffected (Fig. 7biii), in contrast to what was observed in mice^3^ (Extended Data Fig. 10aiii).

**Figure 7.**
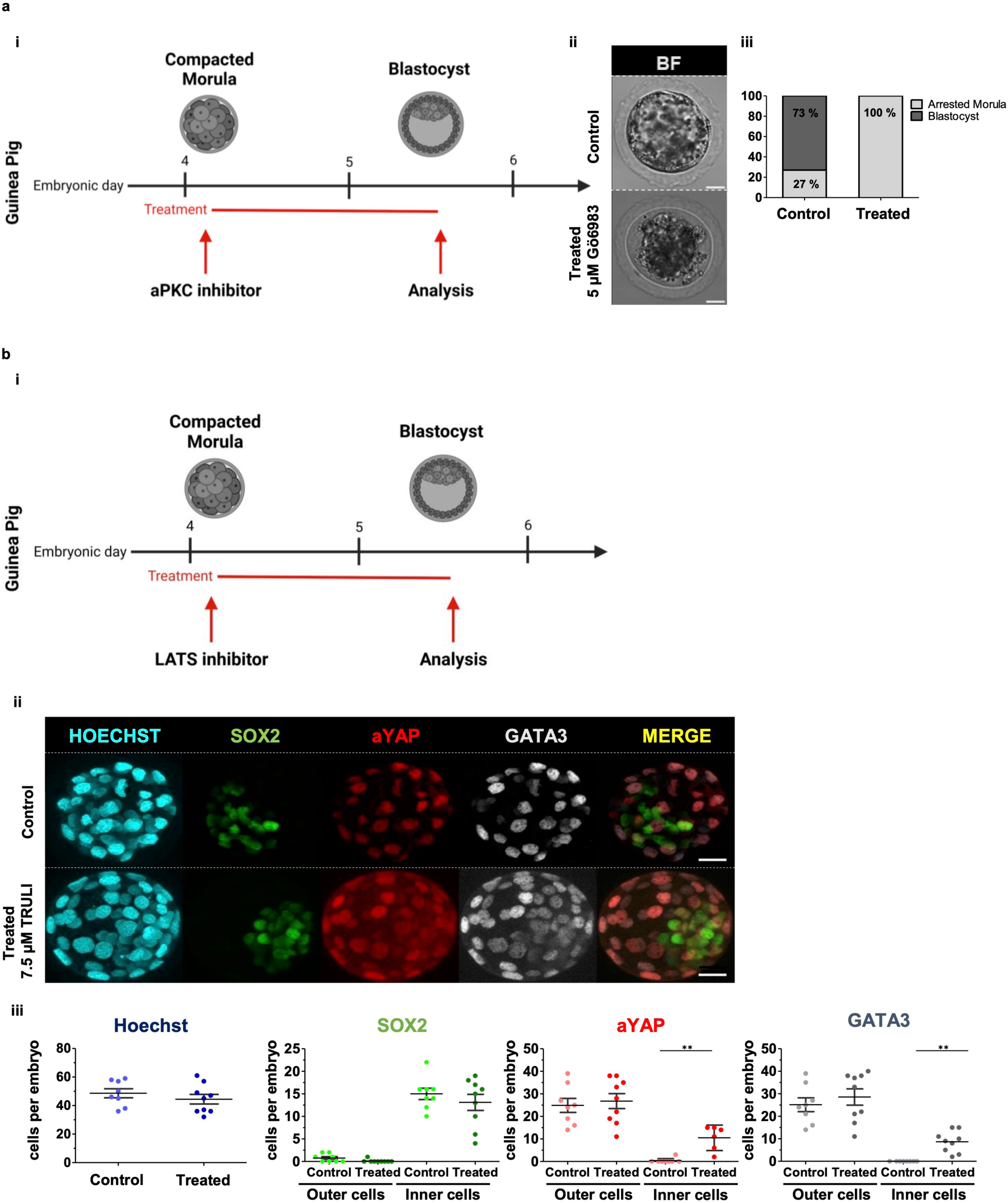
PKC and LATS inhibitor treatments in guinea pig embryos. (a) PKC inhibitor treatment: (i) Schematic protocol used in guinea pig embryos; (ii) bright-field representative images of embryos at E5.5 for control (DMSO) and Gö6983 PKC inhibitor treated guinea pig embryos and (iii) quantification of percentage of guinea pig embryos either developing to form a blastocyst or arrested morula in control (n = 8) and PKC inhibitor treated (n = 3) embryos. (b) LATs inhibitor treatment: (i) Schematic protocol used in guinea pig embryos; (ii) Immunofluorescence analysis of SOX2 (green), aYAP (red), GATA3 (grey) and Hoechst nuclear staining (cyan) in control (DMSO) and LATS inhibitor treated guinea pig embryos and (iii) Scatter plot showing the quantification of the total number of cells per embryo (Hoechst stained) and the number of cells per embryo for the indicated markers in control embryos (n = 8) and embryos treated with LATS inhibitor: 7.5 µM of TRULI (n = 9). For b(ii) ** p ≤ 0.001, Mann–Whitney test. Scatter plots with Mean ± SEM are also indicated in the graph. Scale bars: 20 µm.

To investigate whether the timing of exposure to TRULI (16-cell versus 8-cell in the guinea pig and mouse, respectively) influenced the observed disparities in the effect of Hippo signaling inactivation on Sox2 in inner cells, we conducted an additional experiment. Recognizing that Sox2 expression in mice begins at the 16-cell compacted morula stage, we then treated mouse embryos from the 16-cell stage to the late blastocyst (Extended Data Fig. 10bi-iii). In treated mouse embryos, Lats inhibition led to a significant ectopic nuclear localization of aYAP and GATA3 in inner cells. Notably, the number of outer cells expressing aYAP and GATA3 at late blastocyst in treated embryos did not differ from controls, in contrast to shorter exposure (8-cell to early blastocyst). This suggests that the embryo regulates the number of cells in the TE just before implantation. Upon Lats inhibition, the number of Sox2-expressing cells in inner cells decreased to zero (5/7 embryos) but was not completely abolished in some embryos (2/7) at the late blastocyst stage (Extended Data Fig. 10bii-iii). This implies that in certain cases, a residual number of cells expressing Sox2 can be found in the embryo if the treatment starts later. In summary, Hippo inactivation in guinea pig and mouse embryos promotes ectopic upregulation of TE markers YAP and GATA3 in the ICM, indicating an evolutionarily conserved role for the Hippo signalling pathway mediated through LATS kinases in TE program activation. However, unlike in mice, Hippo inactivation in guinea pig embryo inner cells does not seem to affect SOX2 expression to the same extent.

In mouse and rat preimplantation embryos, the inhibition of Fgf/Mek-Erk pathway has a profound impact on lineage segregation^62–65^ (Fig. 8a). Blocking FGF signalling through MEK-ERK using PD0325901 (PD032) redirects all cells of the ICM to the EPI fate, bypassing the PE. However, PD032 exhibits different effects in other mammals. For instance, in rabbit^66^ and bovine^67^ embryos, PD032 completely abolishes the expression of the PE marker SOX17, suggesting an impairment on the PE formation but it does not increase the proportion of SOX2+ or NANOG+ cells in the ICM compared to control embryos, as seen in mice and rats. In human and porcine embryos, inhibiting MEK-ERK signalling with low dose of PD032 does not abolish the expression of PE markers SOX17 and GATA4, although it reduces the number of SOX17+ cells in porcine embryos^42^ and GATA4+ cells in human embryos^65^. In porcine embryos, a concentration of PD032 (10 µM) diminishes the number of PE cells, resulting in <3 SOX17+ cells per embryo; and there is an apparent, though not significant, shift towards NANOG+ cells in the ICM. The role of FGF/MEK-ERK signaling in the guinea pig has not been explored. However, our scRNA-seq analysis demonstrated an up-regulation of *FGFR2* expression in guinea pig PE cells, suggesting a possible role of FGF in PE formation or expansion. We cultured 16-cell guinea pig embryos with MEK-ERK inhibitor PD0325901 (10 µM)^42,67^, and control until blastocyst stage. We observed a decrease rather than an abolition of SOX17+ cells (Fig. 8bii-iii), as previously reported in the porcine and human PE^42,65^. Of note, the high concentration of PD032 impacts the total number of cells compared to the control. We next wanted to examine the impact on the EPI cells of the guinea pig embryo. Taking this into account, we analyzed the ratio of SOX2+ cells/total number of cells marked by Hoechst. We found that even when the embryo had a lower number of total cells and therefore lower number of SOX2+ cells with the treatment, the ratio of pluripotent cells does not change (Fig 8biii). This is similarly observed in rabbit and bovine embryos and in contrast to mice (Fig. 8 aii-iii), where inhibition of MEK-ERK signaling results in the expansion of the EPI compartment due to all ICM cells converting to pluripotency^66^. In humans, it is difficult to determine the effect on the EPI population due to the high variability within embryos treated with PD032^65^. Moreover, in guinea pigs as in humans, PD032 treatment may affect the viability or proliferation of cells, since the total number of blastocyst cells with treatment is lower than the control. Our findings contribute to the species-specific role of MEK-ERK signaling on early embryonic development, highlighting the intricate interplay between signaling pathways and cell fate determination.

**Figure 8.**
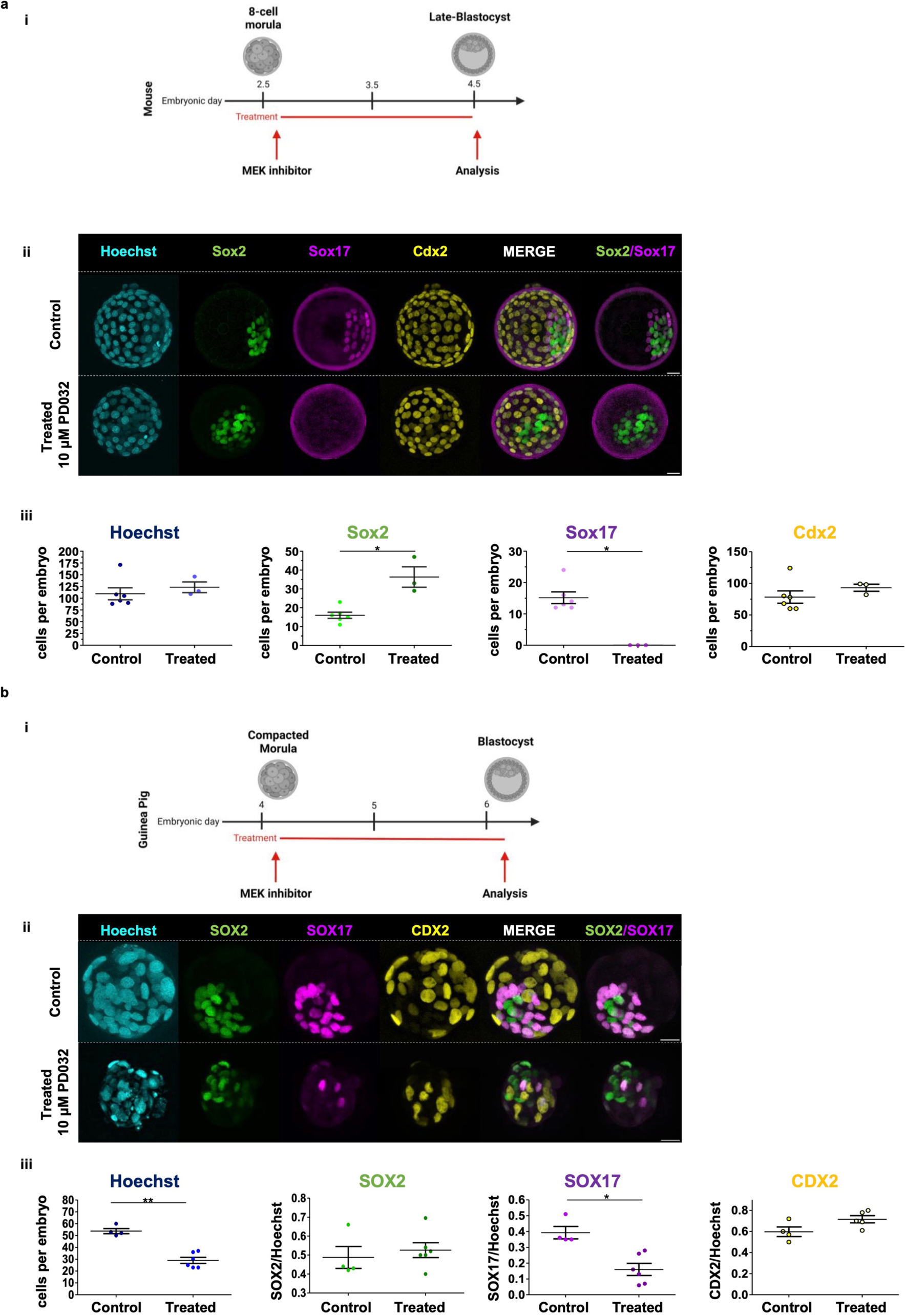
MEK-ERK inhibitor treatment in mouse and guinea pig embryos. (a) MEK-ERK inhibitor treatment in mice: (i) Schematic protocol used in mouse embryos; (ii) Immunofluorescence analysis of Sox2 (green), Sox17 (magenta), Cdx2 (yellow) and Hoechst nuclear staining (cyan) in control (DMSO) and MEK-ERK inhibitor-treated mouse embryos (*p ≤ 0.05, Mann–Whitney test for Sox2 and in a Wilcoxon test for Sox17) and (iii) Scatter plot showing the quantification of the total number of cells per embryo (Hoechst stained) and the number of cells per embryo for the indicated markers in control embryos (n = 6) and embryos treated with MEK-ERK inhibitor: 1 µM of PD0325901 (n = 3). (b) MEK-ERK inhibitor treatment in guinea pigs: (i) Schematic protocol used in guinea pig embryos; (ii) Immunofluorescence analysis of SOX2 (green), SOX17 (magenta), CDX2 (yellow) and Hoechst nuclear staining (cyan) in control (DMSO) and MEK-ERK inhibitor-treated guinea pig embryos (*p ≤ 0.05, Mann– Whitney test) and (iii) Scatter plot showing the quantification of the total number of cells per embryo (Hoechst stained) and the ratio of number of cells for the indicated markers and the total number of cells per embryo in control embryos (n = 4) and embryos treated with MEK-ERK inhibitor: 10 µM of PD0325901 (n = 6). Scatter plots with Mean ± SEM are also indicated in the graph. Scale bars: 20 µm.

### X-Chromosome Activity

In eutherian preimplantation development, X-chromosome inactivation (XCI) involves the silencing of one X-chromosome (X-Chr) in mammalian females (XX) to balance X-Chr output with males (XY). We and others have reported species differences between rodents and primates and as such aimed to examine the status of the two X-Chrs in female blastocysts of the guinea pig. In mice, a paternally imprinted X-Chr is observed starting at the 4-cell stage and maintained in the TE and PE lineages. Following implantation, a reactivation of the X-Chr occurs in the EPI, followed by random maternal or paternal XCI^68^. In humans and human naïve stem cells, we and others have demonstrated a dual dosage compensation in the female (XX) involving the dampening of activity in both X-Chrs^24,69^. First, we performed RNA FISH of the long noncoding RNA *XIST* in both human and mouse male and female embryos (Fig. 9a and Extended Data Fig. 11a). In the mouse preimplantation embryo, we observed one cloud of *XIST* in female embryos, marking the inactivated X-Chr as expected. In the human preimplantation embryo, we replicated our previous results and observed two *XIST* clouds localized to the two X-Chrs (Fig. 9a). Despite multiple attempts, we were unable to perform RNA FISH in guinea pigs. This was likely due to the inability to design sufficient probes given that the genome is predominantly represented as scaffolds, and there are gaps in the assembly. As such, we performed immunostaining, using H3K27me3 as a proxy for XCI (Fig. 9bi-ii). H3K27me3 is an epigenetic mark typically associated with silencing and is found to accumulate on the inactive X-Chr in mouse embryos^70^ and to localize to both X-Chrs in naïve stem cells which exhibited dual X-Chr activity^69^. We first confirmed that H3K27me3 expression mirrored that observed by *XIST* RNA FISH, with one mark observed in the mouse and two in human female blastocysts (Fig. 9 bi-ii). This now provides the first evidence of H3K27me3 localization to the XaXa in human female blastocysts, similar to what has been reported in naive stem cells and fitting with the dual dosage compensation model for X-Chrs^24,71^. Male blastocysts, in contrast, lacked H3K27me3 (Fig. 9bii). Similar to what we observed in the human blastocysts, in E6 guinea pig embryos, we observed two H3K27me3 marks for female blastocysts and no marks in male embryos (Fig. 9bi-ii and Extended Data Fig. 11b). This suggests that X-Chr activity in the female preimplantation guinea pig embryo is similar to the human, where dual dosage compensation occurs, unveiling a new *in vivo* model to better understand the different mechanism(s) underlying XCI in primates.

**Figure 9.**
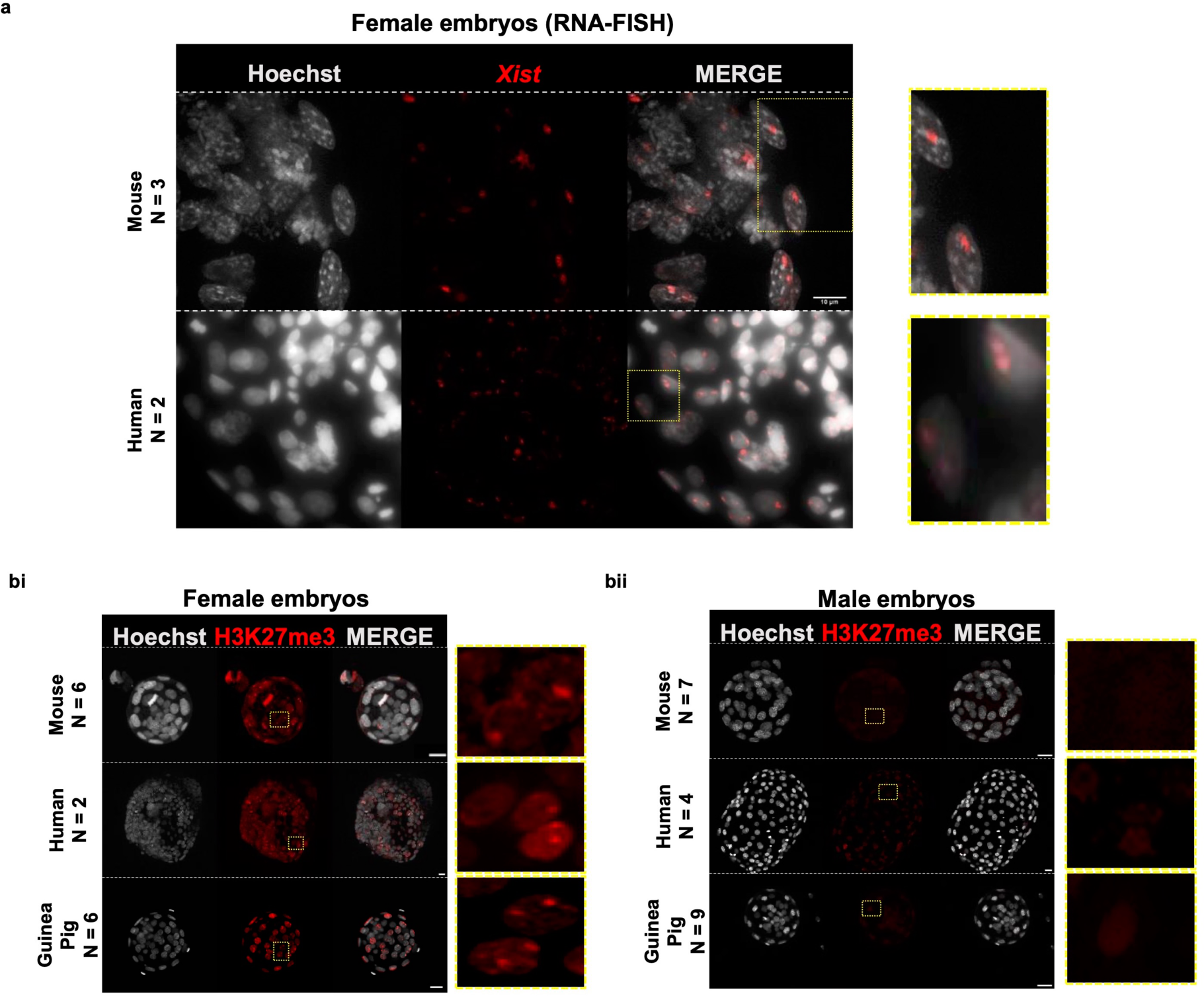
X-Chromosome Status in Mouse, Human, and Guinea Pig Female Blastocysts. (a) RNA FISH detection of *Xist* (red) and Hoechst (gray) in female embryos of mouse (n = 3) and human (n = 2). (b) Immunofluorescence staining of H3K27me3 (red) and Hoechst (gray) in female embryos of mouse (n = 6), human (n = 2) and guinea pig (n = 6). (c) Immunofluorescence staining of H3K27me3 (red) and Hoechst (gray) in male embryos of mouse (n = 7), human (n = 4) and guinea pig (n = 9). Dotted yellow square sections for each embryo show a higher magnification area with the pattern of Xist (a) or the H3K27me3 (b and c) expression. Scale bars: 20 µm.

## DISCUSSION

In this study, we delve into the intricacies of guinea pig blastocyst development, aiming to unveil key molecular features governing early lineage segregation, pluripotency, and X-inactivation. By comparing our findings with established data from human and mouse embryos, we determined both the conserved and divergent features in the early stages of mammalian development. We combined a comprehensive profiling of single-cell transcriptome and protein expression throughout preimplantation as well as functional analysis of conserved pathways, offering insights into the regulatory events shaping the specification of embryonic and extraembryonic lineages in guinea pig embryos and contributing to a broader understanding of the molecular landscape in early mammalian embryogenesis.

We now propose that the guinea pig presents a very powerful small animal model which can be utilized to enhance our understanding of both comparative biology and human embryogenesis. Considering the growing clinical use of vitrification for storing human embryos, obtaining access to very early human embryos is becoming increasingly rare, particularly before E5 when the blastocyst is already formed, and lineages are specified. As such, our ability to study the very early stage of human embryogenesis (zygote to morula) is difficult. Stem cell models of the 4-cell and 8-cell stages still require optimization to more closely recapitulate the totipotent state observed in the human. Though we did not assess zygote or 4-cell embryos, at the 8-cell stage we observe a remarkable similarity in terms of morphology and transcriptome and we now speculate that the guinea pig may serve as a model to better understand this period in humans.

In the guinea pig, the onset of CDX2 expression occurs during the 16-32 cell window, just prior to cavitation in the outer cells, albeit with high co-expression with SOX2, followed by a rapid transition to only CDX2+ outer cells observed around cavitation. This pattern of expression is similar to what has been reported in the human embryo, with co-expression first occurring at E5 followed by a transition to CDX2+ outer cells coinciding with blastocoel formation^20^. Perhaps investigation of human embryos at smaller time intervals (e.g. 2-3hrs) during E5 would demonstrate a similar transition to what we observed in the guinea pig, with a poised expressed of CDX2/SOX2 in outer cells, followed by the initiation of CDX2 only expression in 1-2 cells and then a rapid spreading of CDX2+ only in the outer cells.

In guinea pig embryos, implantation is orchestrated by the mural TE^72^, in contrast to humans where implantation is initiated by the polar TE, adjacent to the epiblast. Successful implantation is a major barrier toward establishing a viable pregnancy. Different strategies for implantation are utilized by mammals including centric (like rabbits, dogs, domestic animals and many marsupials), eccentric (like mouse, rats and hamsters) and interstitial (like humans, guinea pigs and some primates)^11^. Remarkably, primates and the guinea pig (despite being categorized in the Rodentia family), share interstitial implantation, making the guinea pig an excellent model to understand this process with direct translation to the human. We have now for the first time identified a gene signature for polar and mural TE in the guinea pig. Cross-species comparison with the human revealed a remarkable similarity in gene signatures for the opposing TE lineages. In the guinea pig, in contrast to the human, it is the mural cells that proliferate and aid the embryo in undergoing interstitial implantation. Once nearly fully internalised, the mural cells change in appearance with large areas of nuclear-free cytoplasm and some syncytia. At this stage, it is the polar trophoblast that takes over, proliferating under/at the uterine epithelium merging with the mural TE to form the ectoplacental cone^73^. As such, it is not surprising that genes involved predominantly in implantation, would be expressed in the mural TE of the guinea pig, as opposed to the polar as observed in the human. The polar TE in humans (mural in the guinea pig), maintains its stemness (proliferative and preventing TE differentiation) and forms the extraembryonic ectoderm (ExE) of the nascent egg cylinder^74^. Of particular interest, we noted high expression of nuclear receptor subfamily 2, group F, member 2 (NR2F2) marking the mural TE of the guinea pig, a recently reported marker of polar TE in the human embryo^19^, which is not observed in the mouse. NR2F2 is a nuclear hormone receptor demonstrated to be important for cytotrophoblast (CTB) differentiation^47^. Further, retinoic acid-inducible transcription factor TFAP2A (also known as activator protein 2α or AP-2α) is also involved in the regulation of human villous CTB differentiation. TFAP2A expression is induced by retinoic acid^75^ and NR2F2 has been shown to induce CTB differentiation by activating TFAP2A which is potentiated by RARA and RXRA^47^. Indeed, we did identify an enrichment of binding motif for retinoid X receptor (RXR) and TFAP2C in the mural TE of the guinea pig, albeit not at significant levels, suggesting that perhaps NR2F2 may also play a role in CTB differentiation via coordinated action with retinoid acid. Moreover, during implantation, mammals express retinoids and as such, the mural NR2F2 localization in the guinea pig and polar TE expression in the human, may serve as an important receptor for initiating implantation and/or act as a sensor of the implantation window. Similarly, the additional conserved genes identified in the mural (guinea pig) and polar (human) TE, may point to particularly important genes involved in implantation and TE differentiation.

Here we determine that atypical PKC and the evolutionarily conserved Hippo signaling are also involved in blastocyst formation and the establishment of the guinea pig TE. However, similar to what has previously been reported in the human, mouse, rat, and bovine, we also observe that not all the downstream factors are conserved in their expression dynamics.. The role of LATS1/2 in preventing ectopic expression of TE in the ICM of the guinea pig embryo, appears similar to that observed in the human, where a decrease but not complete abolition of SOX2 in the ICM was observed^3^. These downstream differences may be linked to the conserved timing of lineage specification observed between guinea pigs and humans, which differs from mice or perhaps a difference in SOX2 regulation and/or turnover. Further, the timing of TE commitment occurs earlier in the mouse compared with the human^27,57^. When lineage commitment occurs in the guinea pig remains to be determined, however given the striking similarities identified in our study, we speculate that it will occur post-implantation, as observed in humans^57^.

Moreover, MEK-ERK signalling pathway appears to have a similar role in guinea pig as in human PE formation. In mouse^63^, bovine^67^ and rabbit^66^, inhibition of the MEK-ERK pathway completely abolishes SOX17 expression. Whereas in guinea pig, human^65^, marmoset^25^ and pig^42^ embryos, it has a relatively modest impact. Notably, substantial reduction in SOX17 expression requires elevated concentrations of MEK inhibition in guinea pig embryos, which remains to be tested in the human but is consistent with findings in pig embryos^42^. This discrepancy in dosage of MEK-ERK inhibitor suggests that alternative pathways contribute to PE formation mammalian embryogenesis. Moreover, the lack of a clear expansion of the epiblast cells after inhibiting MEK-ERK pathway in humans, bovine, rabbits and guinea pigs suggest the lack of a compensatory interplay between cells of the ICM (EPI/PE) as observed in the mouse. In humans, the window of lineage plasticity is known to extend until after implantation, whereby cells from the TE could convert back to EPI^57^. In mice, we have demonstrated lineage commitment of the ICM and TE occurs at late 16 and early 32 cell stage, respectively^24,27^. It would be interesting to investigate if human, guinea pig and rabbit epiblast cells have a more unrestricted plasticity in the embryo which in turn enables the ability to regulate the number of cells between lineages allowingfor interplay, as observed for the TE and EPI. Overall, we now demonstrate that signaling pathways shown to be conserved in mouse, rat, bovine and humans are also conserved in the guinea pigs embryos. Importantly, these pathways appear to impact the downstream transcription factors and lineage populations of the guinea pig similar to what has been reported in the human, further underscoring the guinea pig as an appropriate model for studying lineage specification in humans.

In placental female mammals, XCI is the silencing of one X-Chr. In mESCs and mouse preimplantation embryos, lncRNA *XIST* is a master regulator of XCI and acts in *cis* to silence the active X-Chr. *XIST* recruits the polycomb group complex PRC2 to the X-Chr that subsequently catalyzes the repressive epigenetic mark, tri-methylates histone 3 on lysine 27 (H3K27me3), which inturn leads to further silencing of the X-Chr^76^. In mouse preimplantation embryos and mESCs, a close relationship has been reported between *XIST* and the status of the X-Chr^77,78^. In contrast, an uncoupling between *XIST* activity and X-Chr status occurs in human blastocysts and naive hPSCs^24,69,79^, which we have previously reported to be associated with bi-allelic dampening. In human embryos, the transition to random XaXi is believed to have initiated around E8 but not fully resolved by E12^80^. The presence of H3K27me3 nuclear foci is often used to identify the inactivated X-Chr in XX female somatic cells. However, recent reports using naive stem cells, which model the preimplantation embryo, have demonstrated the presence of two H3K27me3 loci, co-localized with *XIST* and XaXa^71^. In human preimplantation embryos, the accumulation of H3K27me3 remains controversial, with one report demonstrating H3K27me3 accumulation in 30% of cells by E6^81^. However, in both the rabbit and Cynomolgus monkey, the accumulation of H3K27me3 on the two *XIST*-expressing active X-Chrs has been shown^79,82^. Our study is now the first report demonstrating the accumulation of H3K27me3 and *XIST* around the two female X-Chrs, during which time bi-allelic dosage compensation occurs in the human female blastocyst. The dynamics of XCI in human development are still poorly understood and little is known about the underlying mechanism(s) governing the transition from dampened X-dosage compensation to random XCI in the human post-implantation embryo. Cross-species studies may help us better understand and gain an appreciation of the various mechanism(s) utilized to balance X-Chr output in mammals. Further, employing various models is valuable considering the limitations, in terms of accessibility, ethical, and legal, associated with human embryo research, but one must consider if the chosen model is a good fit for the aspect of development they are trying to understand. Given the resemblance and possible bi-allelic X-Chr activity occurring in the guinea pig preimplantation embryo, we now suggest that the guinea pig may serve as a robust *in vivo* model for assessing both the biallelic compensation and the resolution of the dual dosage compensation observed in humans and shed light onto the mechanism(s) involved. Resolution of the reads obtained from the X-Chr and Y-Chr as well as the exact sequences for *XIST* and *XACT* non-coding RNA can provide further insights into the disconnect between H3K27me3, *XIST* and X-Chr dosage compensation, which at this time is not feasible due to the incomplete guinea pig genome assembly.

As any modeling system, it is crucial to acknowledge both similarities and disparities inherent in what is being modeled. Throughout this manuscript, we have underscored numerous advantages of employing the guinea pig as a model for investigating early embryogenesis and the lasting health impacts on offspring due to exposures or ‘insults.’ In addition, it is important to address limitations of modeling systems. First being that the current guinea pig genome assembly, established 15 years ago, is outdated and accessible only at the scaffold level, resulting in substantial genomic gaps. Moreover, both Ensembl and RefSeq encounter issues such as inaccurate transcription boundaries, missing isoforms, and unannotated genes. While we have taken precautions to ensure accurate gene transcription regions via genome-guided transcriptome assembly, the incomplete genome assembly, coupled with insufficient sequence depth and coverage biases in single-cell RNA-seq data, may still present challenges. Undoubtedly, there is a need for the establishment of a more comprehensive guinea pig genome assembly reference. This advancement will facilitate comprehensive multi-omic studies and cross-species analyses, providing a deeper understanding of evolutionarily conserved and divergent mechanisms underlying embryogenesis. Though currently at the chromosome level, the rabbit genome similarly presents issues with accurate gene annotations and was last updated in 2009 (rabbit genome OryCun2.0)^83^. In addition, current sub-optimal superovulation protocols and the guinea pig’s tendency to produce small litter sizes (typically 2-3 embryos) pose challenges. This limited embryo yield not only increases experimental costs but also constrains the number of manipulations that can be simultaneously conducted, thereby reducing throughput. Furthermore, culturing conditions, particularly for the very early stages (zygote to compaction), are not yet optimized for the guinea pig. This limits *in vitro* manipulation experiments from post-compaction until the blastocyst stage. Our laboratory is actively working to optimize these aspects, aiming to enhance the flexibility and utility of the guinea pig model for studies on early embryogenesis. We have developed an online ShinyApp for exploring guinea pig gene expression (https://petropoulos-lanner-labs.clintec.ki.se/app/ShinyGpigPreImpEM), which we believe will serve as a valuable resource for the scientific community.

Overall, our study demonstrates that the guinea pig embryo during preimplantation development could serve as an ideal small animal model to further interrogate blastocyst formation, lineage specification and possibly XCI, in order to better understand these processes in humans. Data from this study uses a comparative biology approach to gain insight into important evolutionarily conserved and divergent mechanisms driving embryo formation within mammals. We would like to highlight that findings pertaining to embryogenesis simply by Class (mammalia), without considering Order, Family, Genus and Species, may result in inappropriate or erroneous extrapolations to mammalia, emphasizing the need for a more nuanced and species-specific approach. Finally, the guinea pig provides an *in vivo* system that is not under similar ethical and legal constraints which researchers face when working with human embryos and allows for genetic modification of specific genes or transcription factors as well as allowing the assessment of longer-term phenotype associated with exposures during preimplantation development; opening up new avenues for research and holds promise for advancing our understanding of early human development.

## AUTHOR CONTRIBUTIONS

S.P. conceived the study; S.P. supervised the project; J.C. and S.P designed and performed the experiments; C.Z. performed the bioinformatic analysis; J.C., S.P., and C.Z. analyzed the data, interpreted the results and wrote the manuscript.

## DATA ACCESS

All raw and processed sequencing data generated in this study have been submitted to ArrayExpress under accession numbers GSE253670.

## ACKNOWLEDGEMENTS

This work was supported by funding from the Centre de Recherche du Centre Hospitalier de l’Université de Montréal, grants from The Research Center in Reproduction and Fertility (CRRF), Swedish Research Council (2016-01919), Swedish Society for Medical Research (Dnr4-236-2107) and SP holds the Canada Research Chair in Functional Genomics of Reproduction and Development (950-233204). We acknowledge Savana Biondic for help with TRULI dose response studies in the mouse. The computations and data handling were enabled by resources provided by the Swedish National Infrastructure for Computing (SNIC) at Uppsala partially funded by the Swedish Research Council through grant agreement no. 2018-05973. We thank all individuals and couples whose embryo donations have supported this research.

## METHODS

### Ethical Statement

All procedures involving animals were approved by the Comité Institutionnel de Protection des Animaux (CIPA), IP19022SPci. Human embryos were obtained from the Clinique OVO with ethical approval from the regional ethics board (CERSES-20-107-R and 20.126). Informed written consent was obtained from both parents of all couples that donated spare embryos following IVF treatment. No financial compensation was offered for donations.

### *In vivo* Collection of Guinea Pig Embryos

Hartley guinea pigs (Charles River Labs, Laval, Québec) were housed in the Animal Care Facility at the CRCHUM, maintained at 21 °C with 30-70 % humidity under a 12 h light cycle, and fed with Teklad Global Guinea Pig Diet 2040 pellets, water, and hay ad libitum. Embryos at each stage were retrieved from at least 3 separate Hartley guinea pig females (∼500 grams, 2-3 months of age) between days 3.5 and 5.75 after natural mating. We have previously described the detailed procedure of euthanasia, dissection, and flushing^84^. Briefly, embryos were flushed from the oviduct (E3.5) or uterine horns (E4-E5.75) using 5-10 ml of warm M2 medium (MR-015-D, Sigma Aldrich) supplemented with 1 % Penicillin-Streptomycin (15130122, Life Technologies), washed and briefly kept in M2 drops covered with Ovoil ^TM^ (10029, Vitrolife) at the incubator at 5 % O_2_ 5 % CO_2_ at 37.5 °C prior to being fixed in PFA 4 % for 15 min or single cell picking. E6 embryos were collected at E5.75 and incubated in similar conditions for 6 h to reach the desirable embryonic day, as implantation occurs around E6, and obtaining embryos by flushing the uterus was challenging.

### Immunofluorescence

Immunofluorescence was performed on guinea pig embryos as we previously described^84^. Briefly, embryos were fixed in 4 % paraformaldehyde in PBS at room temperature (RT) for 15 min, permeabilized in PBS 0.25 % Triton X-100 at RT for 10 min, and then blocked in PBS with 3 % BSA for 2 h at RT. Samples were incubated with primary antibody O/N at 4 °C in blocking solution. Samples were incubated with secondary antibodies 2h at RT in blocking solution. Washes were performed after primary and secondary antibody incubations 4 x 5 min in PBS. Hoechst (33342, Invitrogen) was used for nuclear staining. All incubations and washes were carried out in a clean well using Nunc 72 well mini trays (CA62409-296, VWR) with 17 µl of each solution. Embryos were mounted in PBS and placed between two coverslips (1.5 thickness) using SecureSeal spacers (Grace Bio-labs). Antibodies used are listed in Supplementary Table 6.

Images were acquired using an Olympus FV1000MPE confocal microscope equipped with an XLUM Plan FL N 20x/1.00 water objective. For excitation, 405 nm (solid-state), 488 nm (Argon laser), 543 nm (solid-state) and 635nm (solid-state) lasers were used for DAPI, Alexa Fluor (AF) 488, AF594 and AF647, respectively. For detection, photomultiplier tubes (PMT) detectors were set as follows: a first SDM490 was positioned in front of the first PMT associated with a BA 430-470 for DAPI detection; then a SDM560 was positioned in front of the second PMT associated with a BA 535-565 for AF488 detection; a SDM640 was positioned in front of the third PMT associated with a BA 560-660 for AF594 detection; and finally, a mirror was positioned in front of the last detector with a BA 655-755 for AF647 detection. All images were acquired sequentially (frame mode) as follows: AF488 and AF647 simultaneously in first, AF594 in second and DAPI at the end of the sequence. Images were acquired in a 512×512 or 800×800 pixel format with zoom at 4μs/pixel speed with a line Kalmann of 3. Z-stack were acquired with a 1µm step size. Images were acquired using the Olympus Fluoview software (v4.2.3.6, Olympus, Japan). For each embryo, z-stacks were analyzed using FIJI software version 1.53c, allowing virtual labeling based on DNA staining (Hoechst staining) for all individual cell nuclei and counting total number of cells. Using this labeling to identify individual cells, each cell in every embryo was then categorized according to relevant phenotypic criteria, without knowledge of the embryo treatment (blind counting). Phenotypic categories included marker expression (e.g., SOX2 or CDX2 positive or negative), protein localization (nuclear or cytoplasmic), and cell position, with cells in contact with the external environment classified as ‘outside,’ and cells surrounded by other cells considered ‘inside’ cells as in ^23^. For all analysis, background was subtracted and a Gaussian filter was applied. To count cells, we created masks for each channel and used the tool to analyse particles to count the number of cells per embryo as previously reported^67^. We visually inspected the masks to account for any error in counting generated by a wrong segmentation of cells. Scatter plots in Figure 1 were done accounting for number of cells per embryo that only express each lineage marker (SOX2: EPI, SOX17 or GATA6 for PE and CDX2 or GATA3 for TE) and the number of cells that co-expressed different combinations of those markers. marker (Sox2, Sox17 and Cdx2) for mouse embryos. Statistics and graphs were done in Graphpad software version 9.2.0.

### *In vitro* culture of Guinea Pig Embryos

Optimal culturing conditions for guinea pig preimplantation embryos remain to be determined. We experimented with media previously reported to be used with the guinea pig^7^, in addition to what is commonly used in different species including M2 and KSOM (mouse)^85^, RDH (rabbit)^66^, mR1ECM (rat)^3^, GTL^86^ and Global medium^3^ (human) and N2B27 (mouse^34^, human^57^, bovine^67^, ovine^87^, porcine^42^ and blastoids^88^). We found (see Supplementary Table 7) that embryos do not progress *in vitro* in any of the media tested prior to compaction (1C – non-compacted 16C). However, we were able to successfully culture embryos from the compacted 16C stage until late/expanded blastocyst with only N2B27. Preimplantation guinea pig embryos were cultured *in vitro*, as we previously described^84^ to perform the functional analysis studies. Briefly, culture dishes were prepared by placing 50 μl drops of N2B27 culture media on 35 mm plastic dishes, subsequently covered with embryo-tested light mineral oil (Ovoil^TM^ or Embryomax) and equilibrated at 37 °C and 5 % CO_2_ for a minimum of 30 minutes prior to embryo culture. Embryos were collected *in vivo* by flushing with M2 manipulation media from E4-4.25 and then passed through multiple wash drops of N2B27 before culturing them together in one drop of N2B27 medium with DMSO or inhibitors (detailed in the functional analysis section).

### Human and Mouse Embryo Culture

E5 vitrified human embryos were thawed using a vitrification thaw kit (Irvine Scientific; 90137-SO) according to manufacturers’ recommendations. Mouse embryos and human embryos were cultured in droplets of pre-conditioned KSOM (Sigma-Aldrich, MR-121-D) and GTL (Vitrolife; 10145) medium, respectively. These cultures were carefully covered with mineral oil (Ovoil, Vitrolife; 10029). Specifically, IVF 35mm dishes (Nunc, Thermo Scientific; 150255) were used for mouse embryo cultures, while 60 mm IVF dishes (Corning C353802 from Thermo Scientific; 150260) for human embryo cultures. Preimplantation embryos were incubated at 37 °C under normoxic (5% CO_2_ and 21 % O_2_) or hypoxic (5% CO_2_ and 5 % O_2_) conditions for mouse and human embryos, respectively.

### Functional Analysis with Small Molecules

To determine if atypical protein kinase C (aPKC) and Hippo signalling pathways are functionally conserved in guinea pig embryos, we performed a side-by-side comparison with the mouse embryos using small molecule inhibitors. The Large Tumor Suppressor Kinase (LATS) inhibitor (TRULI, Enamine; Z730688380)^3,60^ and the PKC inhibitor (Gö6983, Tocris)^57,89^ were initially dissolved in DMSO to create a stock concentration of 100 mM and 50 mM, respectively. These stock solutions were then diluted to the necessary concentrations using pre-equilibrated embryo culture media. A dose-response experiment, with embryos cultured from the compacted morula stage to blastocyst, was conducted across both species for the PKC inhibitor (mouse: 2-5 µM of Gö6983 and guinea pigs: 2.5-10 µM of Gö6983). The optimal concentration of TRULI, a LATS1/2 inhibitor, used for the mouse was determined by a dose-response (Extended Data Fig. 9) and previous work^3^. The optimal concentration of TRULI for guinea pigs was determined from a range of 5-10 µM considering the impact of the inhibitor on embryo viability and phenotypic outcomes, particularly in terms of active YAP, GATA3, and SOX2 expression at E5.5. The optimal concentration of 7.5 µM for TRULI and 5 µM for Gö6983 was utilized in the experiments presented in the main Figure 7 for guinea pig embryos. For the control group, embryos were cultivated in pre-equilibrated media to which 0.1 % DMSO was added, an equivalent concentration to that used with small molecule inhibitors.

Finally, we conducted a comparative analysis examining the inhibition of the MEK-ERK pathway, known for its conserved role in primitive endoderm (PE) formation across various species including mice, rabbits, bovines, pigs and humans. Based on previous studies conducted on bovine and porcine embryos^42,67^ we used PD0325901 at a concentration of 10 µM and DMSO as control during *in vitro* culture from compacted morula to late blastocyst in both mouse and guinea pig embryos.

Immunofluorescence was performed using antibodies SOX2 (EPI), SOX17 (PE) and CDX2 (TE) and embryos were imaged and analysed to quantify the total number of SOX2+, SOX17+, and CDX2+ cells as described above. In Figure 7 (PKC and TRULI functional analysis) the scatter plots show the total number of cells positive for each marker (SOX2, aYAP and GATA3). We also discriminated by localization in the TRULI experiments (inner versus outer cells). In Figure 8 (MEK-ERK inhibition), scatter plots show the total number of cells per embryo (Hoescht staining) for guinea pig and mouse embryos and the ratio between the number of each marker (SOX2, SOX17 and CDX2) and the total number of cells in guinea pig embryos while representing the total number of cells per embryo of each. Statistics and graphs were done in Graphpad software version 9.2.0.

### Single-Cell RNA-sequencing Library Preparation

Female guinea pigs were flushed between E3.5-E5.75 (n = 4 at 8C, n = 7 at 16C, n = 4 at 16-32C, n = 6 at EB, n = 16 at MB and n = 14 at LB) and embryos were briefly placed in M2 drops under mineral oil for downstream processing. Zona pellucida was removed using tyrodes, and embryos were dissociated into single cells with TrypLE Express (12604-013, Gibco, Life Technologies) collected using fine glass capillaries as we previously described^85^. Libraries were prepared using the Smartseq2 as previously described^24,27,90^. The quantity and quality of the cDNA libraries were assessed using an Agilent 2100 BioAnalyzer (Agilent Technologies). Approximately 1 ng of cDNA per cell was transformed into a single-cell library using the Nextera XT DNA Library prep kit (Illumina FC-131-1096) and following the manufacturer’s instructions with slight modifications. In brief, 1 ng of cDNA (in a maximum of 5 µl of sample volume) underwent tagmentation with 2 µl of TD and 1 µl of ATM at 55 °C for 5 min. Upon completion of the program, the reaction was halted by adding 1 µl of 2 % SDS and incubating for 5 min at RT. Amplification was carried out using 3 µl of NMP per sample and adding 1 µl of each dual-indexed (i7 and i5; Illumina) primer. Individual Nextera XT libraries were pooled and then purified using magnetic beads. Indexed libraries were combined for multiplexing (384 samples per lane). The sequencing was performed using the NovoSeq6000 S4 at 150 bp paired-end.

### RNA Fluorescence in situ Hybridization (FISH)

RNA FISH for the long noncoding RNA *Xist*/*XIST* was performed in mouse and human embryos, respectively, following our previously described protocol^91^. In brief, E3.5-3.75 mouse and E7 human embryos were fixed, placed on a silanized glass coverslip, and air-dried for approximately 2 min. Subsequently, embryos were permeabilized with pre-chilled (−20 °C) methanol (Sigma) for 10 min at −20 °C. After air-drying with methanol for 30 min at RT, the embryos underwent heat shock and were hybridized for 2 h at 38.5 °C in a humidity chamber with *XIST* Quasar 570 (125 nM; SMF-2038-1; BioSearch Technologies) in a hybridization buffer. This buffer comprised RNase-free water, 2X SSC, 10 % w/v dextran sulfate (Sigma), 10 % formamide (ThermoFisher Scientific), 2 mg/ml E. coli tRNA (Sigma), 2 mM ribonucleoside vanadyl complex (New England Biolabs), and 2 mg/ml bovine serum albumin (Jackson ImmunoResearch). Following hybridization, samples were washed with 20 % formamide in 2X SSC. Hoescht 33342 (1 µg/ml; ThermoFisher Scientific) was added to the wash buffer during the final wash. Samples were washed again, mounted with Prolong Diamond antifade (ThermoFisher Scientific) and air-dried in the dark at RT for 24 h before imaging.

### Imaging for FISH

Images were acquired using a Zeiss AxioObserver Z1 Yokogawa CSU-X1 spinning disk confocal inverted microscope (Zeiss, Germany). The microscope was equipped with a motorized stage, Piezo objectives and an Evolve EMCDD (512 x 512, 16 bit, 16 µm pixel size) monochrome camera (Photometrics). Image acquisition used an alphaPlan Apo 100X/1.46 Oil DICIII (UV) M27 objective (Zeiss, Germany) resulting in a final pixel size of 133 nm. For DAPI excitation, a 405 nm (solid state) laser was employed, coupled with a double bandpass filter in emission (480/22 + LP530). Similarly, AF568 excitation utilized a 561 nm (solid state) laser coupled with a double bandpass filter in emission (460/30 + 590/30). Z-stack were executed to capture multiple nuclei, employing a 220 nm step size.

### Sex Determination of Guinea Pig Embryos

After imaging H3K27me3, guinea pig embryos were retrieved from the mounted slides, washed in PBS, and transferred with 2.5 µl of PBS to a tube containing 2.5 µl of Extraction Buffer (PicoPLEX WGA kit, final volume 5 µl). Whole DNA extraction and amplification from single guinea pig embryos were conducted using the PicoPLEX WGA kit following the manufacturer’s recommendations. After amplification, samples were quantified using Qubit, and the sex of the embryo was determined through end-point PCR for guinea pig DYS and SRY genes as previously outlined^92^. DYS-F (GTGTTAATGGTGACAGCATCAGC) and DYS-R (GTGCTGTTGGATCTGAAGTGGAGG) were used for detecting the X-Chromosome while SRY-F1 (CCATGATTGCATTTATGGTGTGGTCCCG) and SRY-R1 (GCCTTTTTTCGGCTTCTGTAAGCATTTTCCAC) were used for detecting Y-Chromosome. PCR reactions (25 µl) were carried out using DreamTaq Hot Start Green PCR mix solution (Thermo Fisher Scientific, Waltham, MA), with 0.4 µM of each Dystrophin primer, 0.8 µM of each Sry primer, and up to 20 ng of genomic DNA. The PCR cycling steps comprised: 95 °C for 3 min, followed by 36 cycles at 95 °C for 30 sec, 58 °C for 30 sec, and 72 °C for 30 sec, with a final cycle at 72 °C for 5 min. PCR amplicons were analyzed on 2 % agarose gels and stained with SYBR Safe DNA gel stain (S33102, Invitrogen). The expected amplicon sizes were 212 base pairs (bp) and 135 bp for Dystrophin and Sry gene amplification, respectively. UV gel images were obtained using the GelXDoc from Bio-Rad with automatic exposure.

### Statistical Analysis

All statistical analyses conducted in this study were executed using GraphPad Prism 9.2.0. Detailed information regarding the number of cells or embryos analyzed (n), the specific statistical tests employed, and the corresponding p-values can be found in each figure or its respective legend. The data presented are typically expressed as mean ± standard error (SEM) unless otherwise stated.

### Improving Transcriptome Mapping for Guinea Pig Single-cell RNA-seq Data

For the guinea pig single-cell RNA-Seq data, adapters and low-quality reads were removed using Trimgalore (https://github.com/FelixKrueger/TrimGalore) with default parameters. Subsequently, high-quality reads were aligned to the guinea pig Ensembl reference genome (v.Cavpor.3.0.105, Cavia porcellus, obtained from the Ensembl website)^93^ using the HISAT2 aligner (v 2.2.0)^94^ with default settings. Only uniquely mapped reads were retained for gene expression quantification. Raw read counts were tentatively quantified using featureCounts (1.6.3)^95^ based on Ensembl reference annotation. However, taking this approach, we initially observed low gene expression for well-conserved lineage marker genes like *SOX17* and *CDX2*, and no gene was annotated as *SOX2*, which we knew was inaccurate given the IF stainings. When we then switched to RefSeq annotation (downloaded from USCS, CarPor3), were we found no expression for *BMP2* and *SOX17*. Visualizing read coverage across the genome, we discovered that a large majority of sequencing reads were aligning to nearby regions and as such incorrectly not aligning to the gene location itself based on the incomplete Ensembl and RefSeq annotations (see Extended Data Fig. 12). We therefore decided to update the guinea pig gene annotation using the following steps: 1. Ensembl transcripts belonging to “lincRNA”, “processed_pseudogene”, “protein_coding”, and “pseudogenes” were selected for the genome guided assembly. The mapping bam files were used as input for Stringtie (v1.3.3b)^96^ for each individual sample. After the initial assembly, the transcripts were merged using ‘stringtie --merge’ and compared with the guided annotation using ‘gffcompare -r’^97^. Transcripts that overlapped with multiple reference genes were excluded, as well as transcripts with a length less than 200 bp and ‘class codes’ identified by gffcompare equal to ‘e’, ‘s’, ‘x’ were removed in downstream analysis. Transcripts with class codes ‘j’, ‘k’, ‘=’, ‘m’, ‘n’, ‘o’, ‘p’, ‘y’ were assigned to corresponding reference genes. Novel transcripts (class codes belong to “u” or “i”) with no clear reference annotation were blasted^98^ against human and mouse transcripts of protein coding and lncRNA with parameters ‘-max_target_seqs 1 -outfmt 6 -num_threads 25 -evalue 1e-20 -strand plus -max_hsps 1’ to find the most similar human or mouse reference genes. If there were no orthologous genes of matched reference genes in the guinea pig Ensembl annotation, they were added. Other novel transcripts with known orthologous genes in the guinea pig Ensembl annotation were reassigned to known annotated genes based on whether the novel transcripts fell within the nearby loci (± 3 kb) of the current annotation. Following this,the transcript and gene relationships were updated in the gene annotation results from gffcompare^97^. Gene annotations (such as miRNA, misc_RNA, Mt_rRNA, Mt_tRNA, ribozyme, rRNA, scaRNA, scoRNA, snRNA, sRNA) that were not included in the genome-guided assembly were included based on their Ensembl annotations in the final reference genome annotation for gene expression quantification using featureCounts with parameters ‘-p -C -D 5000’. Using this genome guided approach, we were then able to successfully identify the previously missing *SOX2* gene, which was absent in the Ensembl annotation. We also accurately identified the 3’ boundary of *NANOG* (ENSCPOG00000008888), *CDX2* and *BMP2*, which were previously misannotated. Additionally, a new isoform (MSTRG.64950.1) of *SOX17*, which was not annotated in Ensembl and RefSeq, was discovered. Together these improvements made a noticeable impact on our downstream analysis.

### Quality Control, Normalization, and Dimensional Reduction Analysis for Guinea Pig single-cell RNA-seq Data

Following that optimization of the transcriptome mapping, low-quality cells were removed based on three criteria, resulting in 541 cells remaining out of the 661 initially sequenced cells. First, each cell was required to express more than 3,000 genes. Second, the expression proportion of mitochondrial genes had to be less than 5 %. Third, the expression proportion of genes related to stress granules^99^ needed to be less than 8.5 % (see Extended Data Fig. 2b). Following the filtering process, single-cell RNAseq expression data were analyzed using the standard Seurat (v4.2.0)^100^ pipeline. Specifically, genes expressed in at least 3 cells were log-normalized for downstream analysis, and the 2000 most variable genes were identified using the "vst" method with the “FindVariableFeatures” function. After assessing the variance explained by technical factors (see Extended Data Fig. 3a) with scater (v1.18.3)^101^ the variance attributed to the ‘number of expressed genes’ was regressed out using the “ScaleData” function. The top 15 principal components were then calculated using the “RunPCA” function and utilized for uniform manifold approximation and projection (UMAP) dimensional reduction. Clusters were identified using the “RunUMAP” and “FindClusters” functions, respectively. Cluster stability was analyzed with R package clustree (v0.4.3)^102^ on a range of resolution values (0.4 to 1.4), with 0.6 yielding the most stable set of clusters. Cell identities were inferred based on previously published lineage markers and cell stages^19^. Marker genes were identified using the “FindAllMarkers” function with default parameters, setting the adjusted p-value cutoff at less than 0.05. Cells belonging to the 8-cell, 16-cell, and precavitation cells from 16-32 cell group were combined as ‘prelineage,’ while ICM and EPI cells were grouped together for marker gene expression analysis. Dimensional reduction for TE sub-lineages were constructed using only the TE cells and top1500 variable genes and top 20 principal components (PCs).

### Profiling Lineage-specific Expression Modules

Self-organizing maps (SOM) were generated using the R package kohonen (v.3.0.12)^35,36^ with the log-transformed expression data from Seurat as input. The grid was set to a 30 × 30 matrix with hexagonal topology. Subsequently, hexagonal grids with a Z-score higher than 1.5 were identified as lineage-specific grids. Functional enrichment analysis was conducted on the genes associated with these grids using the clusterProfiler package (v3.18.1)^103^. KEGG pathway annotations for the guinea pig were obtained from the Ensembl database and downloaded using the “keggList” function from the R package KEGGREST (v1.26.1)^104^.

### Trajectory Interference and Pseudotime Analysis

For the initial analysis to determine the timing of the ICM/EPI and TE segregation, single-cell trajectory analysis was conducted using the R Monocle2 package (v.2.14.0) with the DDRTree method and default parameters^105^ using prelineage, ICM/EPI and TE cells. Specifically, raw counts were used as input for Monocle2 normalization. We selected 450 top differentially expressed genes pre-identified by the “FindAllMarkers” function among clusters to construct the single-cell trajectory and calculating pseudotime by “reduceDimension” and “orderCells” functions from Monocle2. To determine the timing of the second lineage specification (EPI and PE), we performed analysis on the ICM trajectory by only selecting the ICM/EPI, PE, and precavitation cells. Using the Seurat pipeline we generated two-dimensional UMAP based on the top 1500 variable genes and top 20 principal components (PCs). The data was then converted into a Monocle3 (v1.0.0)^106^ object using the “as.cell_data_set” function from the R SeuratWrappers package(v0.3.0) (https://github.com/satijalab/seurat-wrappers). These cells were further categorized into 7 subclusters using the “cluster_cells” function with the “louvain” method. Single-cell trajectory and pseudotime were further inferred using the “learn_graph” and “order_cells” functions from Monocle3, with the precavitation cells having the lowest UMAP_2 values designated as the root cell.

### Differential Expression Analysis and Gene Set Co-regulation Analysis

Differential gene expression analysis was conducted for EPI vs PE, EPI vs TE, and mural vs polar using the "wilcox" test implemented in the FindMarkers function from the Seurat package. Genes with a False Discovery Rate (FDR) less than 0.05, a log2(fold change) greater than 0.25, and expression in more than 10 % of the cells were considered differentially expressed. The contribution of each KEGG pathway to the expression variability of the whole cells was assessed using the “geseca” function with parameters ‘center = FALSE, eps=1e-100’ from the R package fgsea (v.1.28.0)^107^. Significantly variable pathways were identified by requiring a p-value less than 0.05. Z-scores of gene set co-regulation for each pathway were evaluated with the “addGesecaScores” function, using the Seurat object and genes related to the KEGG pathway as input. Gene set enrichment analysis (GSEA) of Differentially Expressed Genes (DEGs) was performed using the “GSEA” function from the clusterProfiler package (v.3.18.1) ^103^.

### Investigation of Motifs in Promoter Regions of DEGs

Position weight matrices of transcription factor binding profiles for vertebrates were obtained from JASPAR2020^108^. Sequences within ± 2 kb of the transcription start sites of DEGs between mural and polar cells were extracted. Subsequently, the transcription factor binding motif enrichment was assessed using the “calcBinnedMotifEnrR” function implemented in the R package monaLisa^109^. Specifically, genes were categorized based on the direction of regulation (up-regulated or down-regulated). Motifs with a p-value less than 0.05 were selected and visualized using the “plotMotifHeatmaps” function."

### Reprocessing single-cell RNA-seq Primate and Mouse Preimplantation Embryo Datasets

Human preimplantation embryo transcriptomes were compiled from two single-cell profiling studies^24,49^. These data were reprocessed as reported in^86^. Updated annotations from Meistermann et al.^19^ were used for the Petropoulos et al. datasets^24,27^. Embryos with lineage segregation at E5 were designated as ‘Early blastocysts’, while those at E6 and E7 were designated as ‘Middle blastocysts’ and ‘Late blastocysts’, respectively. Preimplantation marmoset embryo scRNA-seq transcriptomes were selected from studies by Boroviak et al., 2018^25^, and Bergmann et al., 2022^110^. Expression matrices from both projects were extracted from (https://github.com/Boroviak-Lab/SpatialModelling). The EPI and PE cells in ‘Late Blastocysts’ from Boroviak et al., 2018, were further identified through clustering and expression of EPI and PE known markers as reported in their paper. TE cells from Bergmann et al., 2022, were designated as ‘Middle-Late Blastocyst TE’. Preimplantation cynomolgus monkey embryo scRNA-seq transcriptomes were extracted from Nakamura et al., 2016 ^111^. The data from Nakamura et al., 2016, were reprocessed and remapped using STAR (v2.5.1b)^112^ with the reference genome ‘Macaca_fascicularis_5.0.96’ from Ensembl. Gene expression was quantified using RSEM (V1.3.0)^113^. Previously published annotations downloaded from their publication were used. Embryos belonging to ‘E6’ and ‘E7’ were designated as ‘Early blastocysts’, while those at ‘E8’ and ‘E9’ were designated as ‘Middle or Late Blastocysts’. Preimplantation mouse embryo datasets were selected from two earlier studies, including Deng et al., 2014^50^, and Nowotschin et al., 2019^33^. Data from Deng et al., 2014, were reprocessed and remapped using STAR and RSEM. Expression matrices from Nowotschin et al., 2019, were extracted from the original publication. Since blastocyst lineage information was not included in Deng et al., 2014 only cells belonging to ‘8 cell’ and ‘16 cell’ stages were selected. Cells from Nowotschin et al., belonging to ‘E3.5’, were labeled as ‘Early blastocysts’, while those at ‘E4.5’ were labeled as ‘Middle or Late Blastocysts’.

### Integration of Human and Guinea Pig Preimplantation single-cell RNA-seq Data

To align the human and guinea pig genes, we first converted their gene names to Ensembl gene IDs. We then aligned their Ensembl gene IDs based on the Ensembl ortholog information. Only genes with the ortholog type "ortholog_one2one" were used for integration. After aligning gene features, preimplantation data from human ^24,49^ and guinea pig embryos were integrated together using the canonical correlation analysis (CCA) approach implemented in the R package Seurat. Anchors between the two species were identified using the “FindIntegrationAnchors” function with the top 750 features selected by the “SelectIntegrationFeatures” function. Subsequently, the data were further integrated using the “IntegrateData” function with the following parameters: “k.anchor = 5; k.score = 30; k.weight = 50; k.filter = 100;”. After integration, principal component analysis (PCA) was performed on the integrated data, followed by embedding into UMAP space based on the top 20 PCA dimensions. To align the developmental stages and lineages, mutual nearest neighbors (MNN) between the guinea pig and human preimplantation embryos were identified using the “findMutualNN” function from the R package batchelor (v1.6.2)^114^ on the scaled data with a setting of “k=20”. Pearson correlation between human and guinea pig cells identified as mutual nearest neighbors was calculated on scaled data from the integrated object.

### Comparison of Human, Mouse, and Guinea Pig Lineage Markers

Lineage markers for human mouse and guinea pig were identified using the same “FindAllMarker” function. The union gene set for human, mouse, and guinea pig lineage markers was used to assess the differences among these species. Shared lineage markers were identified for genes with the same gene names expressed in 10 % of the cells and displaying the highest expression in the same corresponding lineages across species.

## Extended Data Figures legends

**Extended Data Fig. 1.**
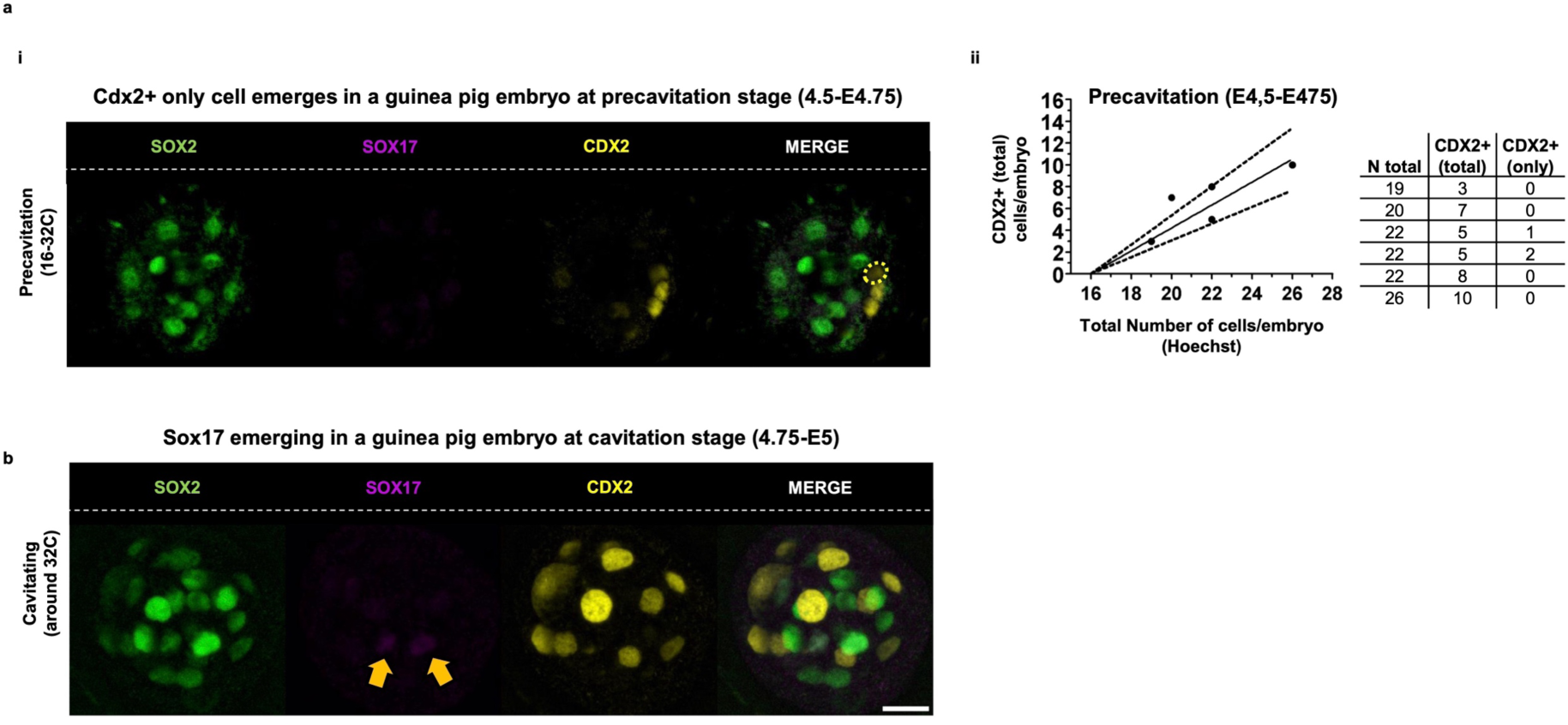
Emerging of CDX2 and SOX17 in guinea pig *in vivo* embryos. (a) CDX2 expression in precavitation stage: (i) Immunofluorescence analysis of EPI (SOX2), PE (SOX17), and TE (CDX2) associated markers in a representative guinea pig embryo during precavitation. The dotted yellow circle indicates an emerging cell with CDX2+ expression only. Only 2/6 embryos presented this at precavitation (ii) Scatter plot depicting the total number of CDX2+ cells per embryo relative to the total number of cells per embryo (Hoechst) (n = 6 embryos collected at E4.5-4.75), along with its linear regression illustrating the variability in CDX2 expression at this stage. (b) SOX17 emerging expression in cavitation: Immunofluorescence analysis of EPI (SOX2), PE (SOX17), and TE (CDX2) associated markers in one guinea pig embryo during cavitation (E4.75-5). The yellow arrows indicate an emerging cell with SOX17+ expression. Only 2 of 4 embryos presented this at cavitation. Scale bar: 20 µm.

**Extended Data Fig. 2.**
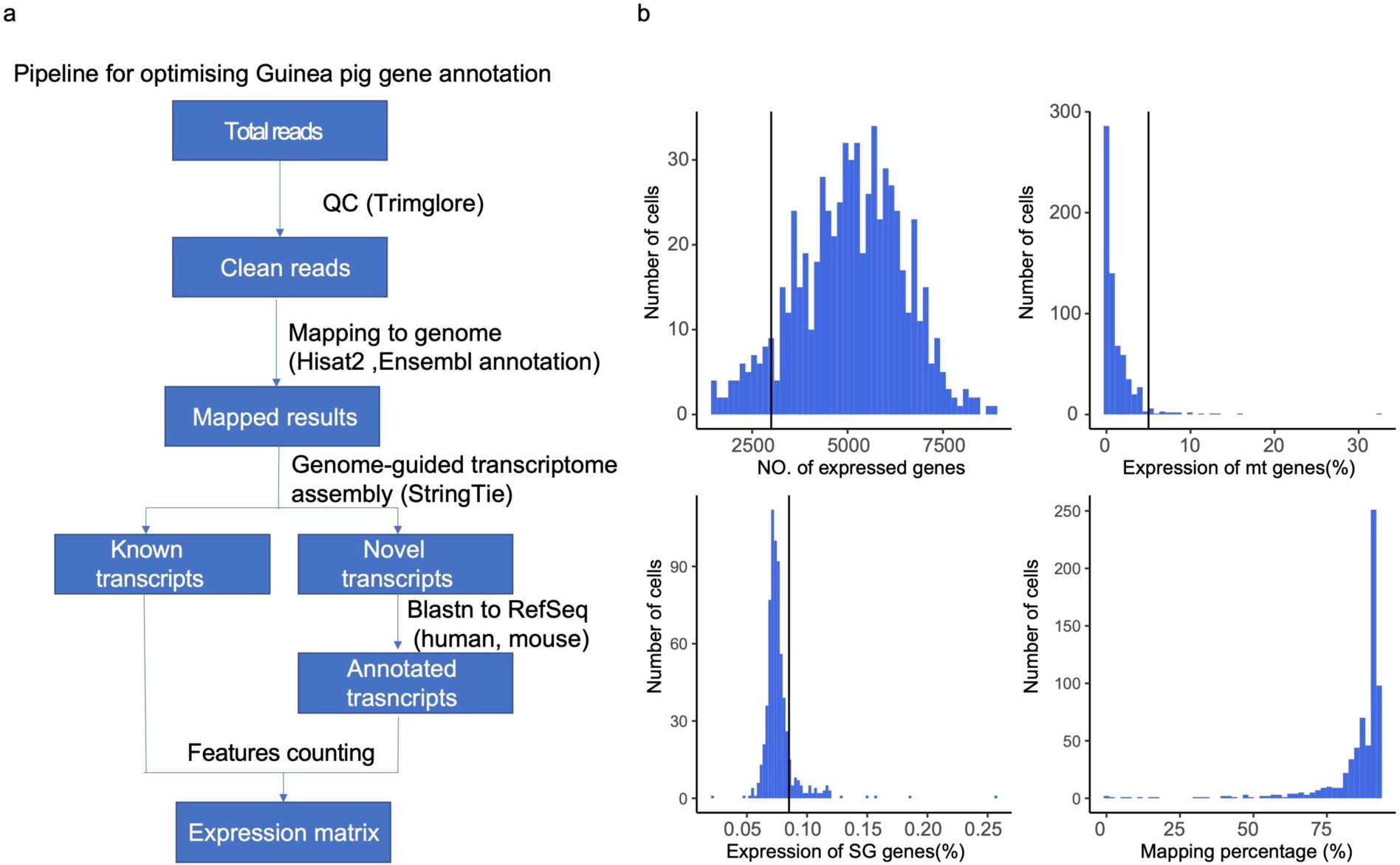
Analysis pipeline and quality control used for guinea pig scRNA-seq dataset. (a) Schematic diagram of optimized guinea pig gene annotation. (b) Histograms display the number of expressed genes, the proportion of mitochondrial(mt) gene expression, the proportion of stress granule(SG) gene expression, and the total mapping percentage. Quality control cutoffs are indicated by black lines.

**Extended Data Fig. 3.**
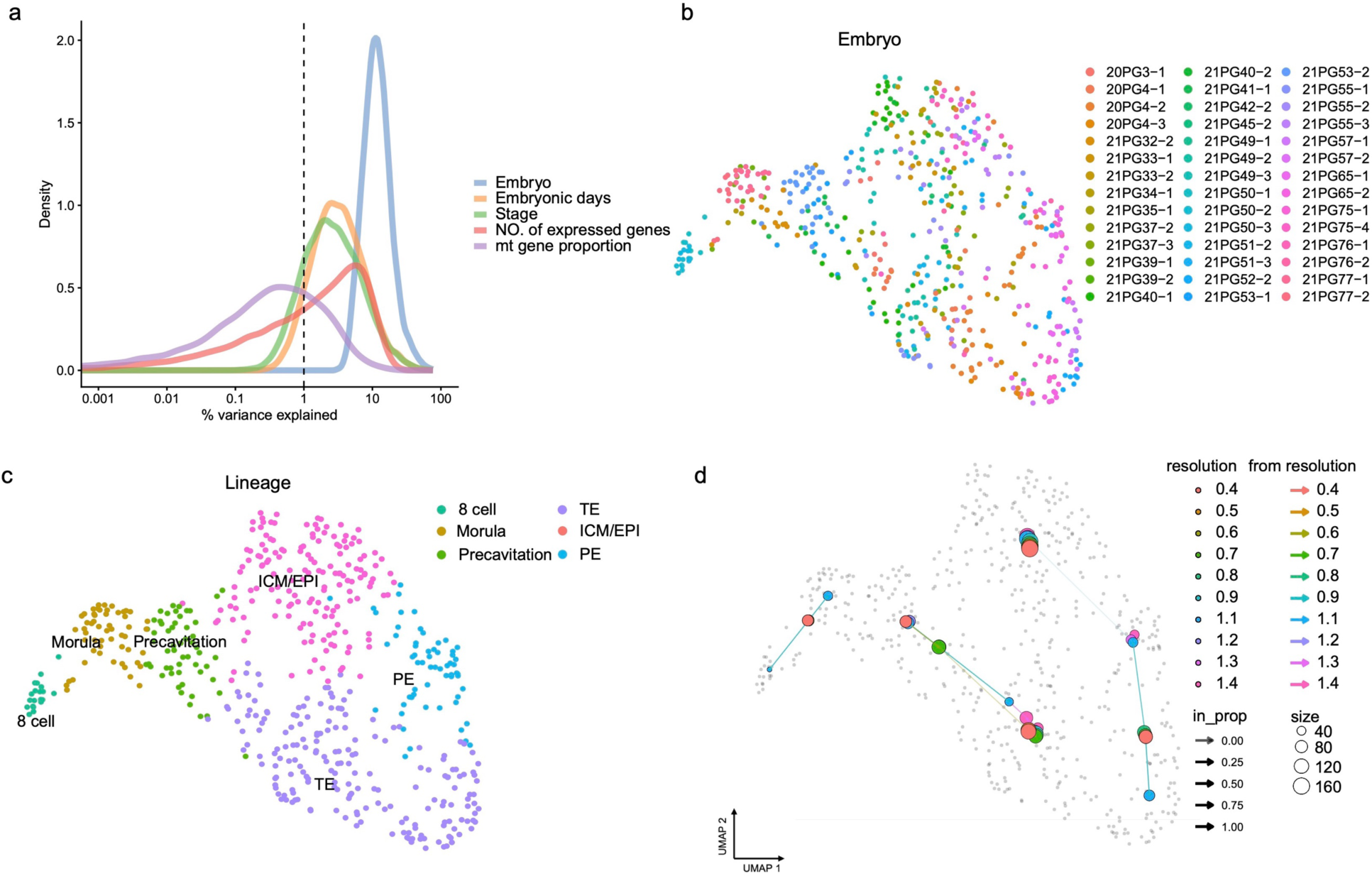
Top variation and resolution exploring for guinea pig scRNA-seq dataset. (a) Line plot showing the percentage of variance explained by factors "Embryo", “Embryonic days”, “Stages", “Number of expressed genes”, and "Mitochondrial gene proportion". (b, c) Two-dimensional UMAP representations of 541 single-cell transcriptomes from guinea pig preimplantation embryos, with colors indicating the embryo source (b) and lineages (c). (d) Cluster stability was analyzed with Clustree at various resolution values (0.4 to 1.4).

**Extended Data Fig.4.**
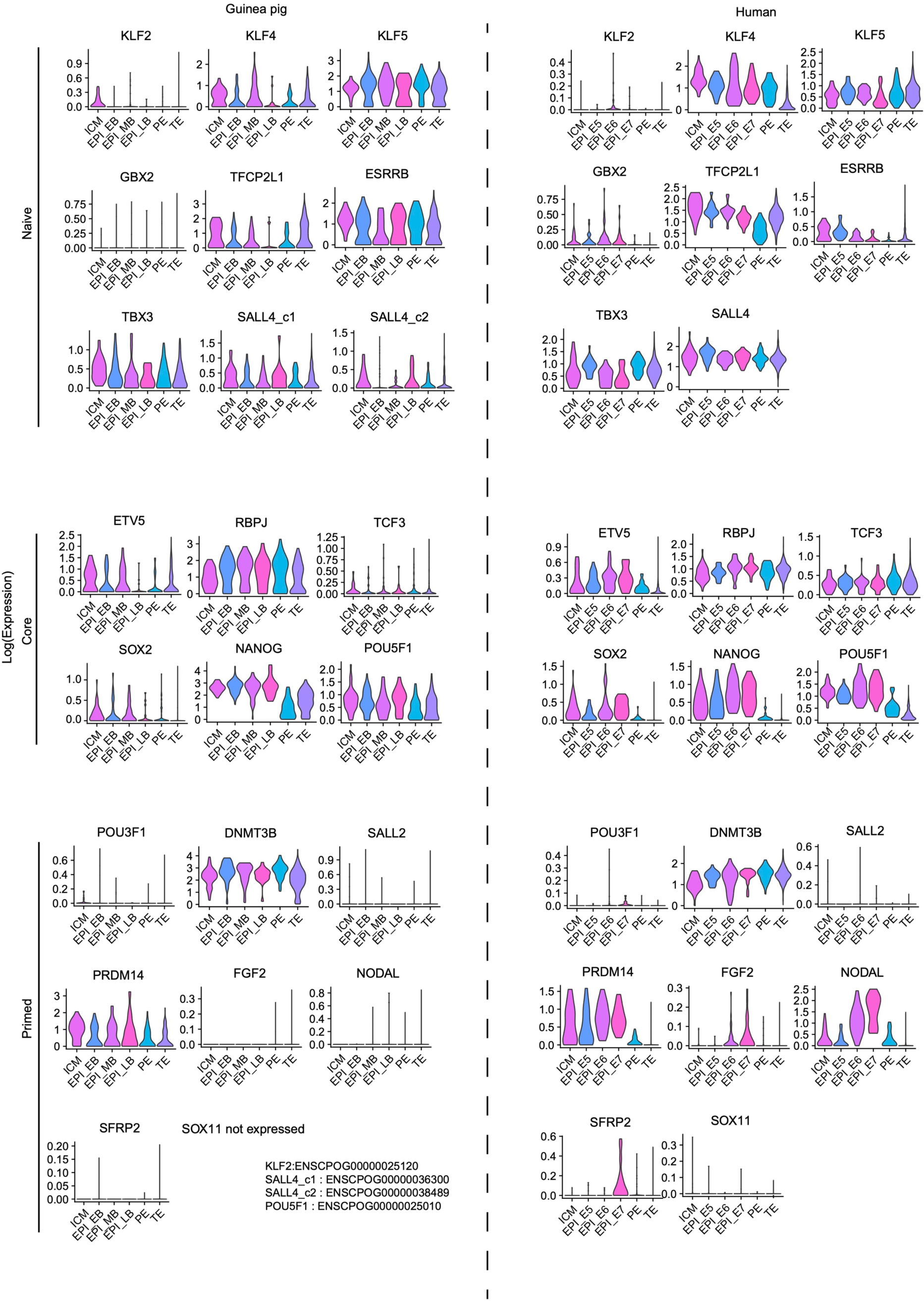
Pluripotency signatures in guinea pig and human preimplantation embryos. Violin plots showing the expression of selected pluripotency (naive, core and primed) genes in guinea pigs and humans, stratified by lineages.

**Extended Data Fig. 5.**
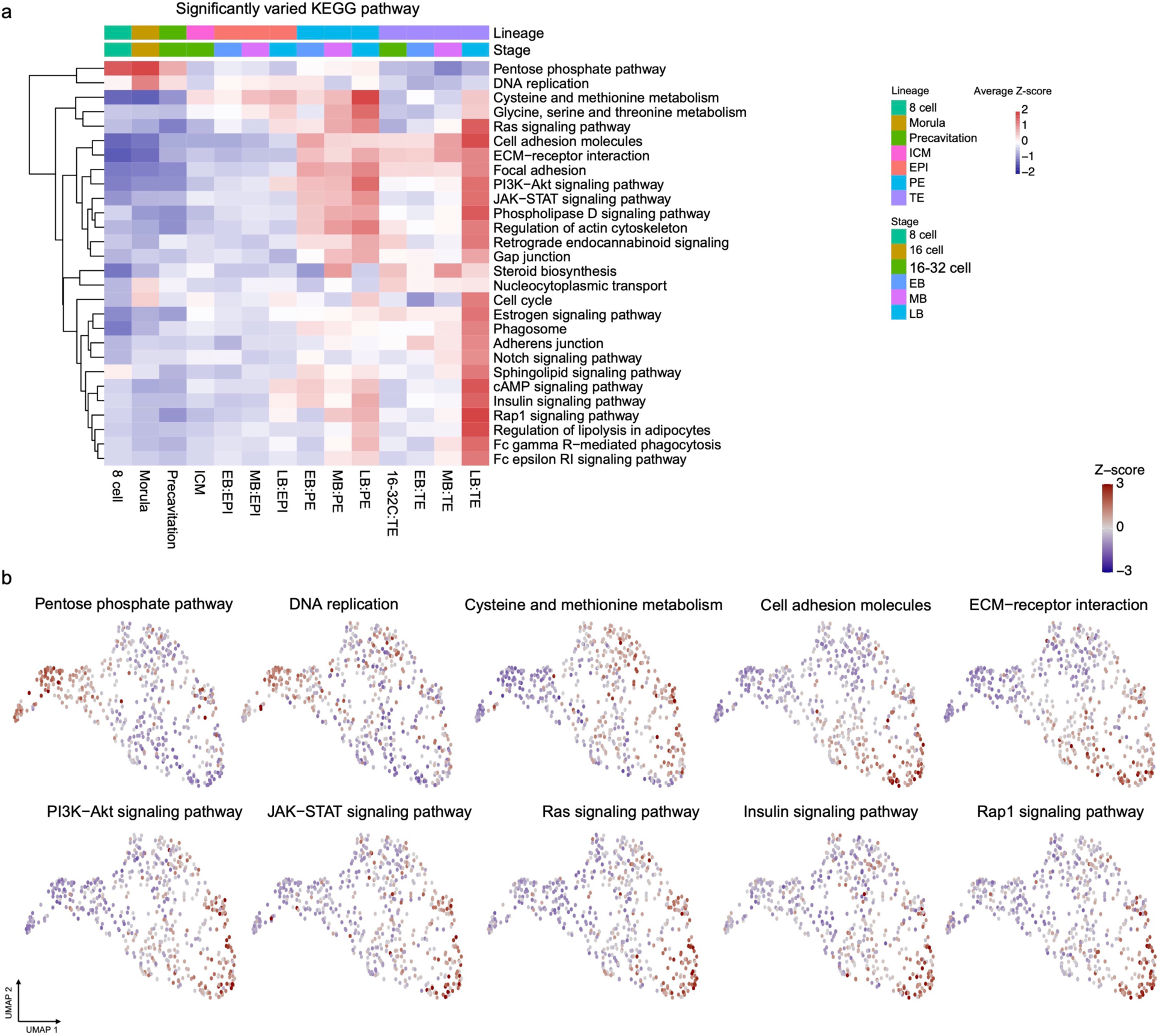
Top variable KEGG pathway in guinea pig scRNA-seq dataset. (a) Heatmap displaying the average Z-score of genes belonging to significantly varied KEGG pathways across all cells, categorized by stages and lineages. KEGG pathways associated with cancer or diseases were excluded from the heatmap. (b) UMAP feature plot of cells exhibiting the average Z-score of genes belonging to selected significantly varied KEGG pathways.

**Extended Data Fig. 6.**
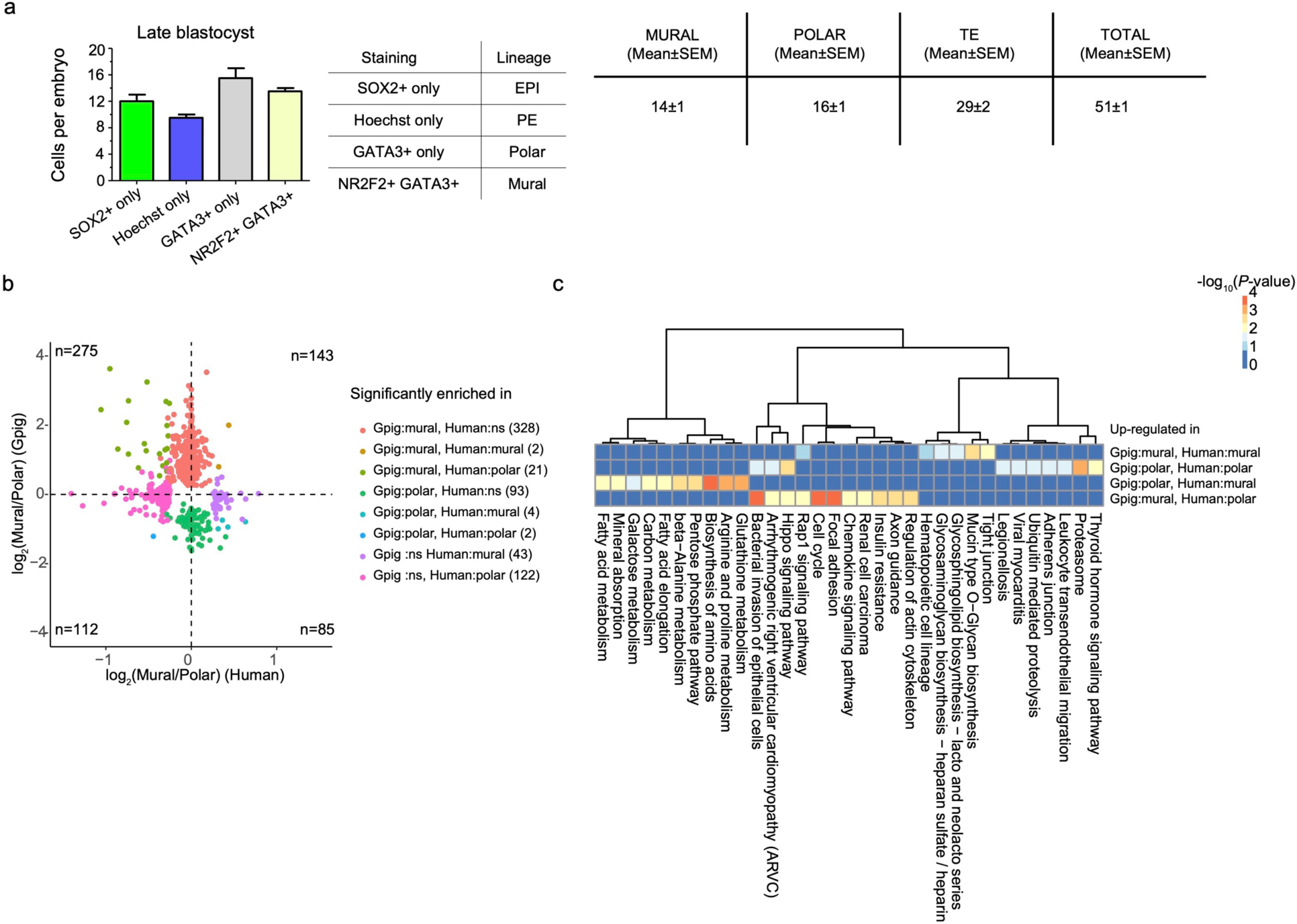
Expression of mural and polar DEGs in guinea pig and human mural and polar cells. (a) Quantification of cells per embryo expressing SOX2+ (EPI), NR2F2+GATA3+ (mural TE) and GATA3+ (polar TE) in guinea pig *in vivo* late blastocysts. Cells with only Hoechst staining represent the PE. Bar graphs represent the Mean ± SEM of n = 2 replicates. (b) Log2 fold change between mural and polar cells in human and guinea pig (x-axis and y-axis, respectively). Colors indicate the differential expression status in the guinea pig and human. The number of genes is indicated behind the color legend. "n" in each corner represents the total number of genes in each quadrant. "ns" denotes "not significant." (c) Heatmap illustrating the significance of enriched KEGG pathways for genes associated with differential expression status in guinea pig and human mural and polar cells.

**Extended Data Fig. 7.**
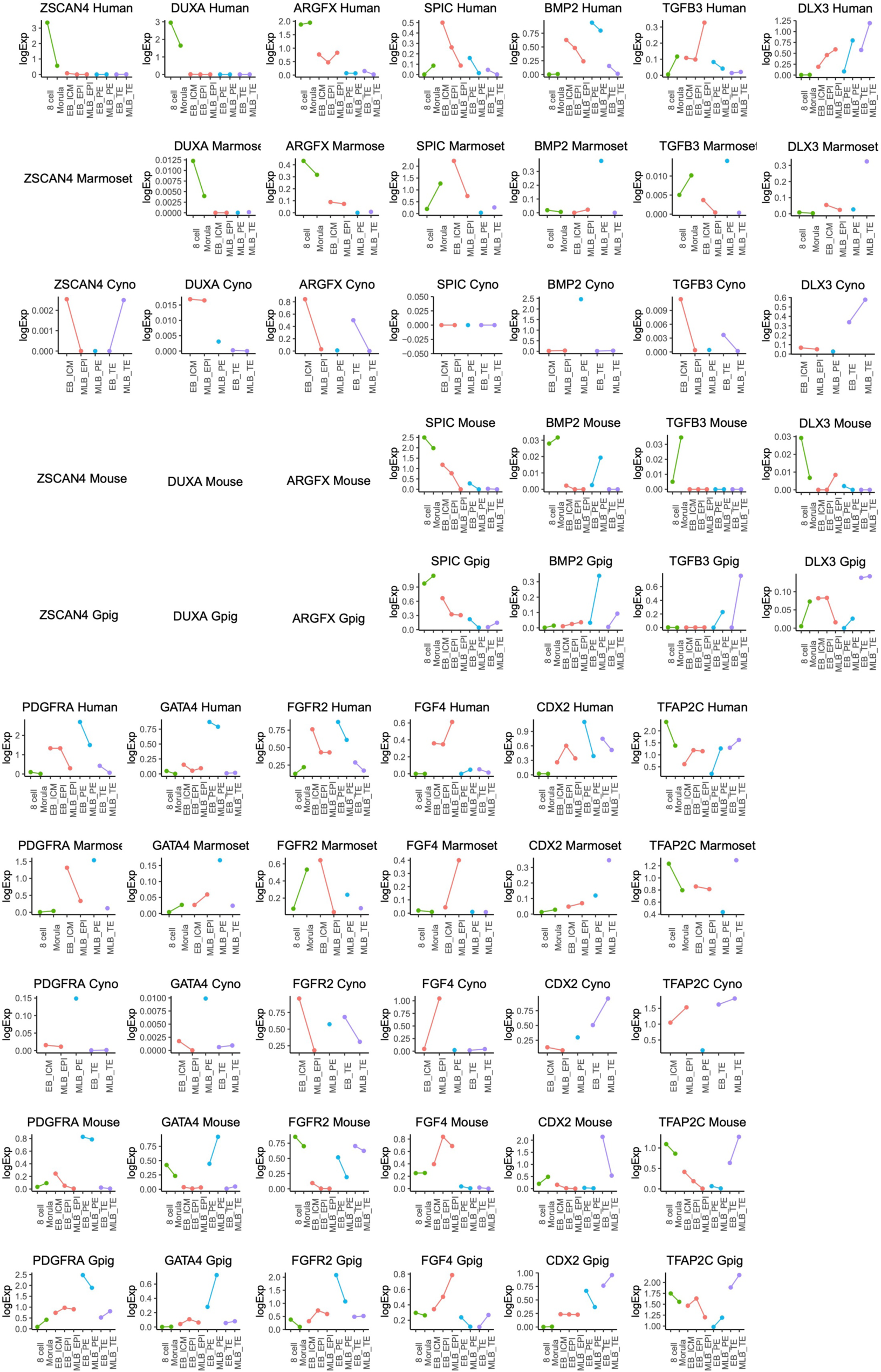
Line plots depict the expression of selected genes, categorized by stages and lineages, across human, marmoset, cynomolgus monkey, guinea pig, and mouse. Prelineage, EPI, PE, and TE are labeled by colors. Genes that could not be detected in the corresponding species are represented as empty.

**Extended Data Fig. 8.**
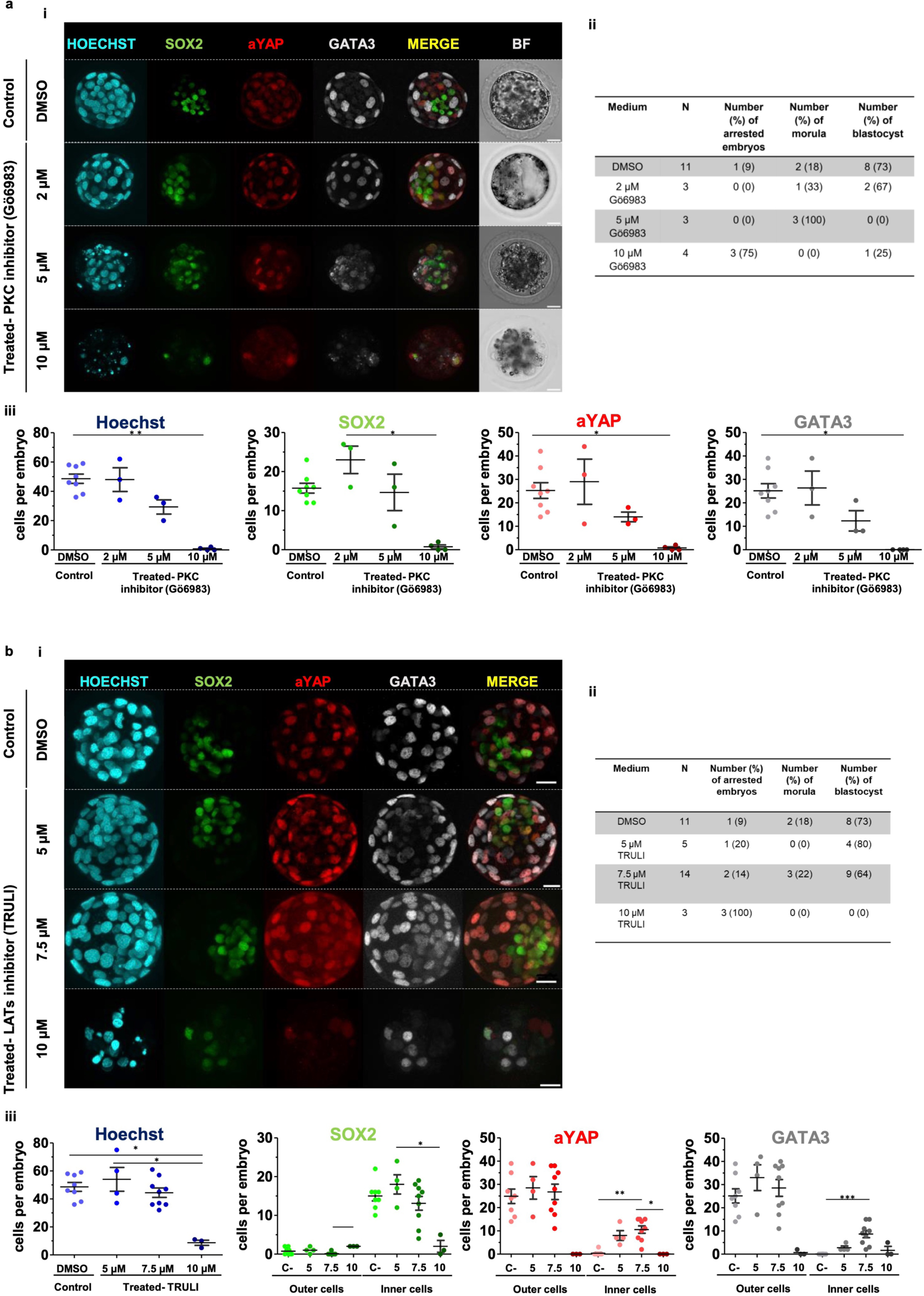
Dose-response of PKCs and LATS inhibitor treatments in guinea pig embryos. (a) Dose-response of PKCs inhibitor Gö6983: (i) Immunofluorescence analysis of SOX2 (green), aYAP (red), GATA3 (gray) and Hoechst nuclear staining (cyan) in control (DMSO) and PKCi inhibitor-treated guinea pig embryos from 16-cell compacted morula to mid-blastocyst. (ii) Quantification of the percentage of guinea pig embryos either developing to form a blastocyst or arrested morula in control and PKCs-inhibitor-treated embryos. (iii) Scatter plot showing the quantification of the total number of cells per embryo (Hoechst stained) and the number of cells per embryo for the indicated markers in control embryos (n = 11) and embryos treated with PKCs-inhibitor: 2 µM (n= 3), 5 µM (n = 3) and 10 µM (n = 4). * p ≤ 0.05 and ** p ≤ 0.01, Mann–Whitney test. Scatter plots with Mean ± SEM. (b) Dose response of LATs inhibitor TRULI: (i) Immunofluorescence analysis of SOX2 (green), aYAP (red), GATA3 (gray) and Hoechst nuclear staining (cyan) in control (DMSO) and LATS inhibitor-treated mouse embryos from 16-cell compacted to morula to late blastocysts. (ii) Quantification of the percentage of guinea pig embryos either developing to form a blastocyst or arrested morula in control and LATs-inhibitor-treated embryos. ** p ≤ 0.05,** p ≤ 0.01, ***p ≤ 0.001 Mann–Whitney test. Scatter plots with Mean ± SEM. (iii) Scatter plot showing the quantification of the total number of cells per embryo (Hoechst stained) and the number of cells per embryo for the indicated markers in control embryos (n = 11) and embryos treated with LATS-inhibitor: 5 µM (n = 5), 7.5 µM (n = 14) and 10 µM (n = 3). Scale bars: 20 µm.

**Extended Data Fig. 9.**
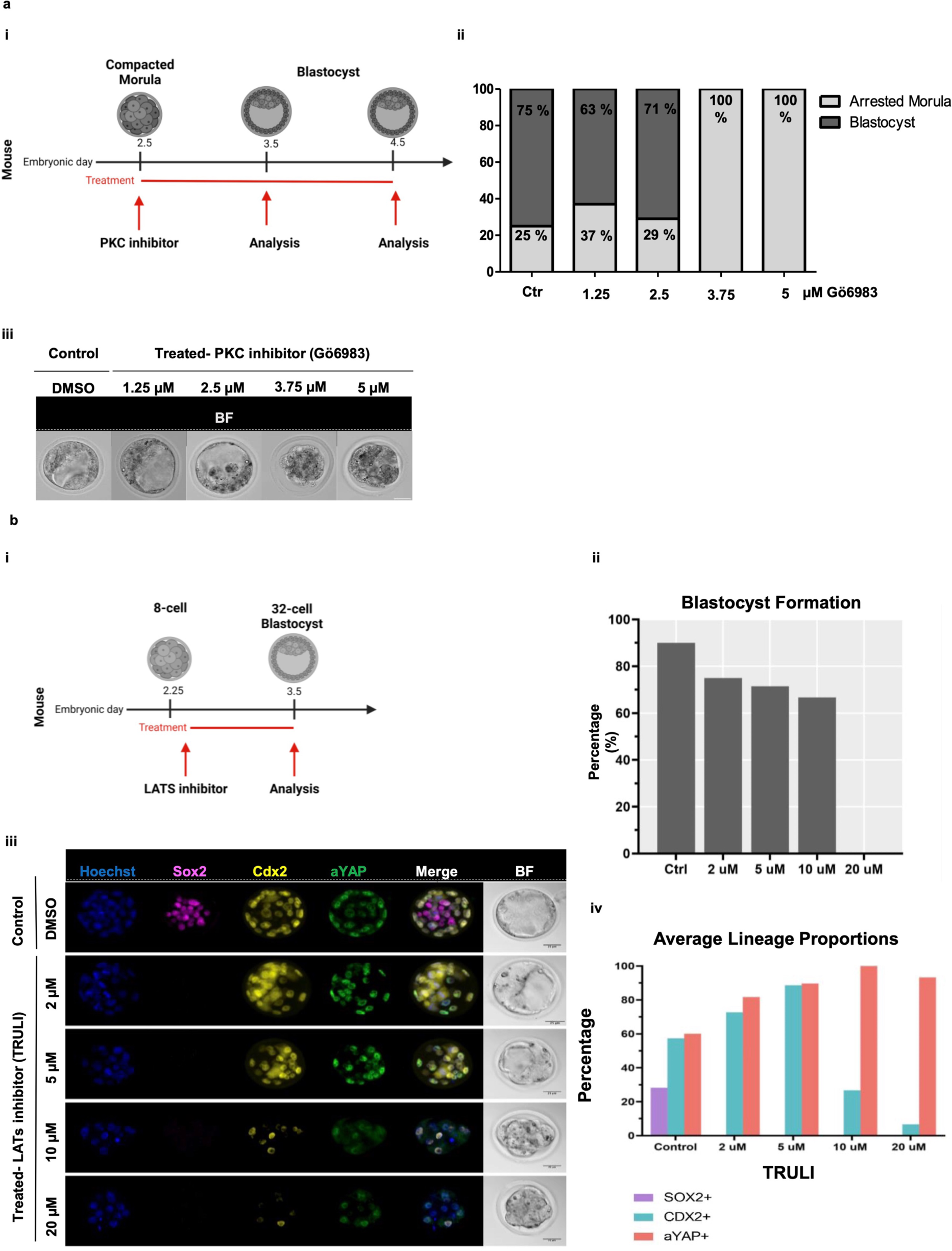
Dose-response of PKC inhibitor and TRULI inhibitor in mouse embryos. (a) Dose-response of PKC inhibitor Gö6983: (i) Schematic of PKC treatments in mouse embryos. (ii) Quantification of the percentage of mouse embryos either developing to form a blastocyst or arrested morula in control and PKCs-inhibitor-treated embryos (1.25-5 µM). (iii) Bright field images of each group. (b) Dose-response of TRULI: (i) Schematic of TRULI treatments in mouse embryos. (ii) Bar plot showing the percentage of embryos developing to blastocyst. (iii) Immunofluorescence analysis of Sox2 (pink), Cdx2 (yellow), aYAP (green) and Hoechst nuclear staining (blue) in control (DMSO) and LATs inhibitor-treated mouse embryos with different concentrations of TRULI inhibitor (2-20 µM) from 8 cell E2.25 to blastocyst E3.5 (n = 3-10 embryos/group). (iv) Bar plot showing the average of lineage proportions. Scale bars: 25 µm.

**Extended Data Fig. 10.**
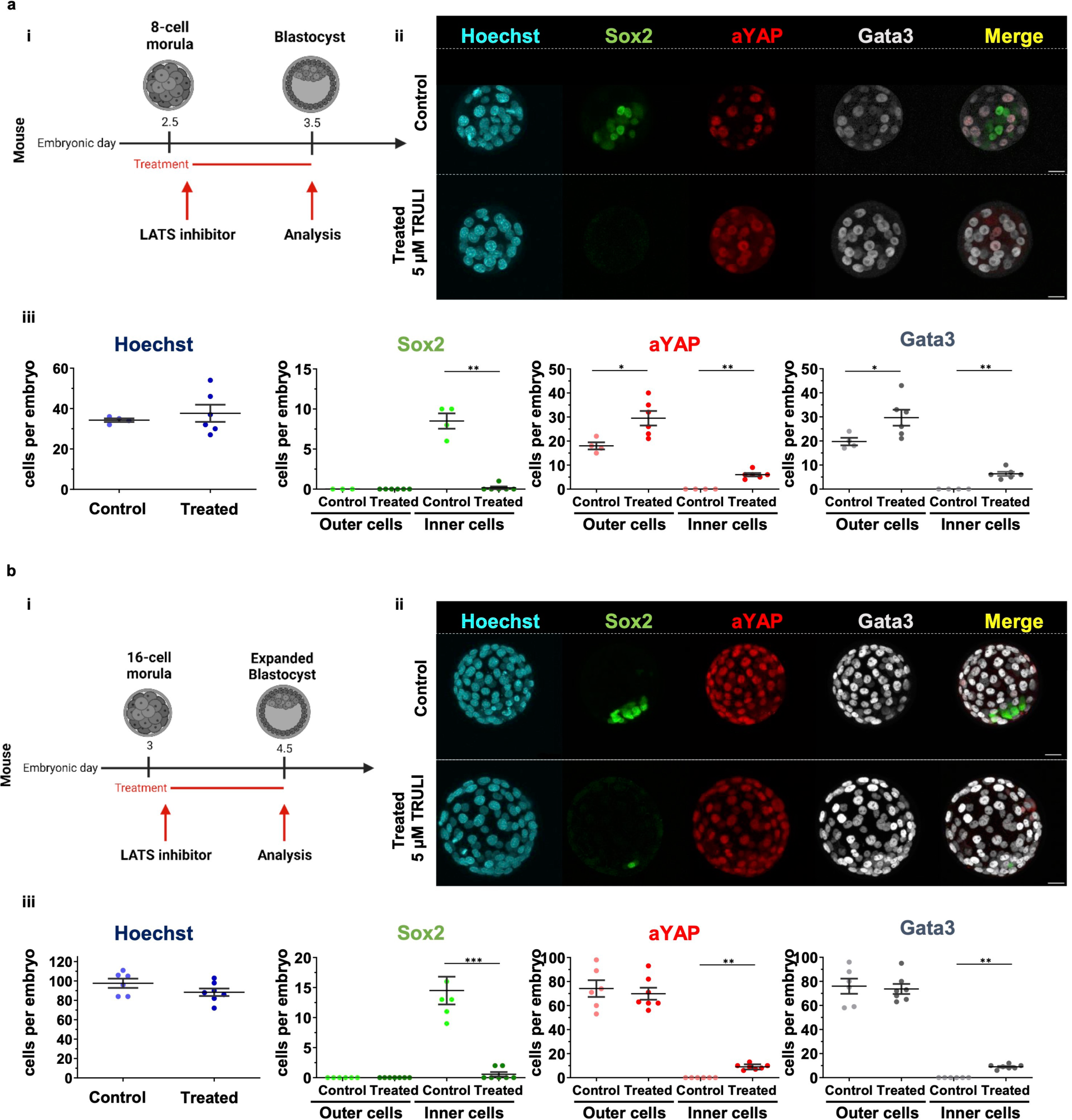
LATS inhibitor treatments in mouse embryos at different starting points. (a) Treatment from 8-cell compacted morula E2.5: (i) Schematic of LATS inhibitor treatment in mouse embryos. (ii) Immunofluorescence analysis of Sox2 (green), aYAP (red), Gata3 (gray) and Hoechst nuclear staining (cyan) in control (DMSO) and LATS inhibitor-treated mouse embryos from 8-cell compacted morula to mid-blastocyst. (iii) Scatter plot showing the quantification of the total number of cells per embryo (Hoechst stained) and the number of cells per embryo for the indicated markers in control embryos (n = 4) and embryos treated with LATS-inhibitor: 5 µM of TRULI (n = 6). * p ≤ 0.05 and ** p ≤ 0.01, Mann–Whitney test. (b) Treatment from 16-cell compacted morula E3: (i) Schematic of LATS-inhibitor treatment in mouse embryos (ii) Immunofluorescence analysis of Sox2 (green), aYAP (red), Gata3 (gray) and Hoechst nuclear staining (cyan) in control (DMSO) and LATS inhibitor-treated mouse embryos from 16-cell compacted morula to late blastocyst. (iii) Scatter plot showing the quantification of the total number of cells per embryo (Hoechst stained) and the number of cells per embryo for the indicated markers in control embryos (n = 3) and embryos treated with LATS-inhibitor: 5 µM of TRULI (n = 3). * p ≤ 0.05 and ** p ≤ 0.01, Mann–Whitney test. Scatter plots with Mean ± SEM. Scale bars: 20 µm.

**Extended Data Fig. 11.**
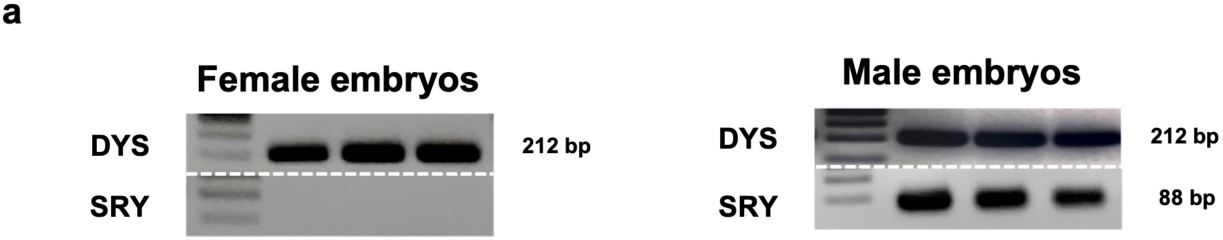
Sex validation of guinea pig embryos following H3K27me3 immunofluorescence. (a) Agarose gel representative image of three female and three male embryos after PCR of DYS and SRY genes of the genomic samples. Amplicon of DYS is 212 bp and of SRY 88 bp.

**Extended Data Fig. 12.**
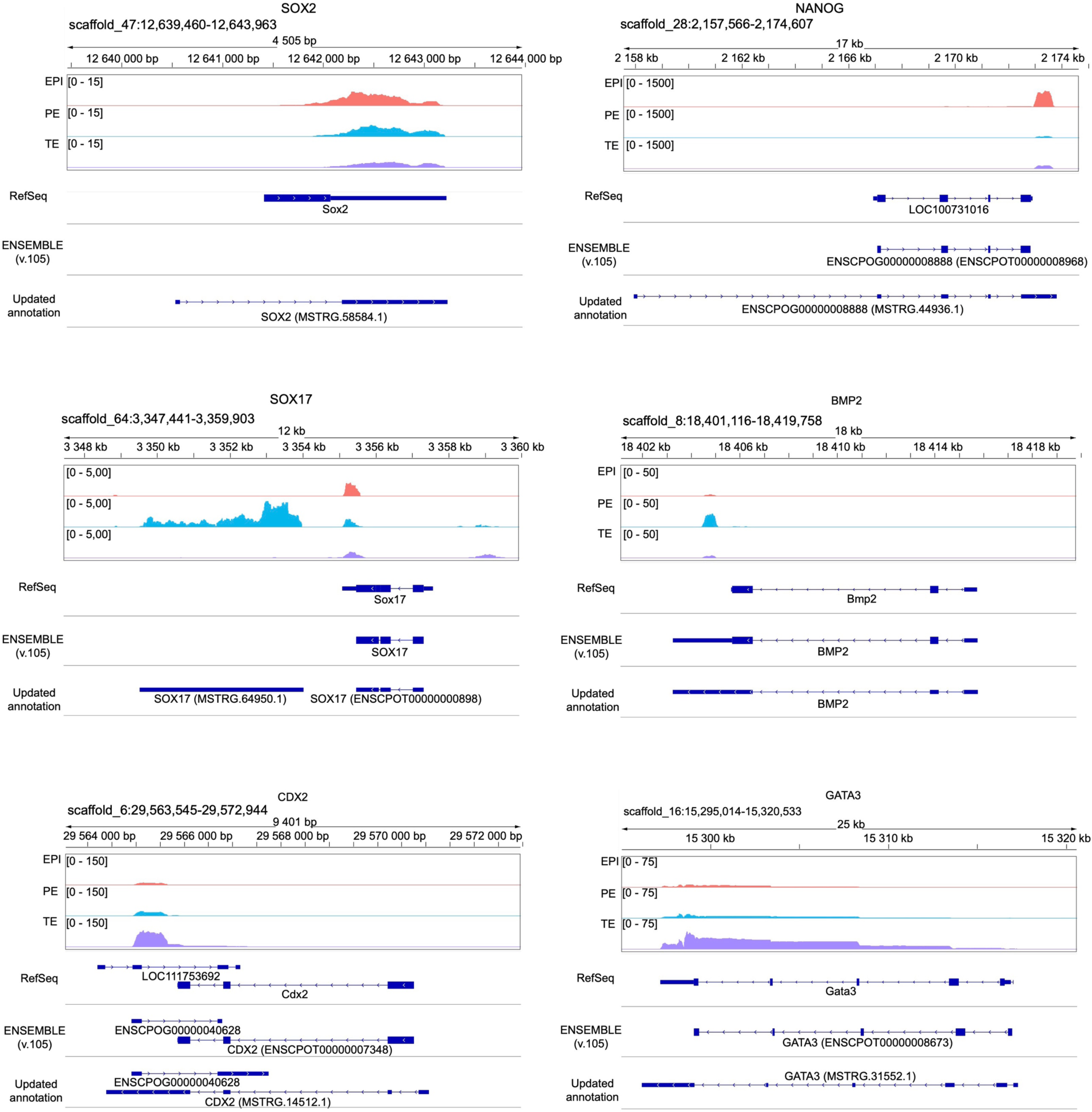
IGV snapshot displaying read distribution from middle and late blastocyst EPI, PE, and TE cells over genes SOX2, NANOG, SOX17, BMP2, CDX2, and GATA3. Gene annotations from RefSeq, ENSEMBL, and updated annotations are listed behind.

## Supplementary Tables titles

**Supplementary Table 1: Meta table for cells that have passed scRNA-seq quality control.**

**Supplementary Table 2 : Lineage markers genes for guinea pig preimplantation.**

**Supplementary Table 3: Table displaying the SOM classification of tissue-specific genes and enriched KEGG pathways.**

**Supplementary Table 4: Differentially expressed genes and enriched pathways between mural and polar TE cells in guinea pig and human.**

**Supplementary Table 5: Table showing identified lineage markers, their average expression, and species conservation in human, mouse, and guinea pig.**

**Supplementary Table 6: Primary antibodies used in guinea pigs embryos. Supplementary Table 7: *In vitro* culture conditions tested for Guinea pig embryos.**

## References

1. Biondic, S., Canizo, J., Vandal, K., Zhao, C. & Petropoulos, S. Cross-species comparison of mouse and human preimplantation development with an emphasis on lineage specification. Reproduction 165, R103–R116 (2023).

2. Gerri, C., Menchero, S., Mahadevaiah, S. K., Turner, J. M. A. & Niakan, K. K. Human Embryogenesis: A Comparative Perspective. Annu. Rev. Cell Dev. Biol. 36, 411–440 (2020).

3. Gerri, C. et al. A conserved role of the Hippo signalling pathway in initiation of the first lineage specification event across mammals. Development 150, (2023).

4. Bouchereau, W. et al. Major transcriptomic, epigenetic and metabolic changes underlie the pluripotency continuum in rabbit preimplantation embryos. Development 149, (2022).

5. Wagner, J. E. Chapter 1 - Introduction and Taxonomy. in The Biology of the Guinea Pig (eds. Wagner, J. E. & Manning, P. J.) 1–4 (Academic Press, 1976).

6. Suzuki, O. et al. Optimization of superovulation induction by human menopausal gonadotropin in guinea pigs based on follicular waves and FSH-receptor homologies. Mol. Reprod. Dev. 64, 219–225 (2003).

7. Suzuki, O. et al. Development of preimplantation guinea-pig embryos in serum-free media. Reprod. Fertil. Dev. 5, 425–432 (1993).

8. Dorsch, M. M., Glage, S. & Hedrich, H. J. Collection and cryopreservation of preimplantation embryos of Cavia porcellus. Lab. Anim. 42, 489–494 (2008).

9. Hribal, R., Guenther, A., Rübensam, K. & Jewgenow, K. Blastocyst recovery and multifactorial gene expression analysis in the wild guinea pig (Cavia aperea). Theriogenology 86, 1299–1307 (2016).

10. Morrison, J. L. et al. Guinea pig models for translation of the developmental origins of health and disease hypothesis into the clinic. J. Physiol. 596, 5535–5569 (2018).

11. Lee, K. Y. & DeMayo, F. J. Animal models of implantation. Reproduction 128, 679–695 (2004).

12. Pfeffer, P. L. Alternative mammalian strategies leading towards gastrulation: losing polar trophoblast (Rauber’s layer) or gaining an epiblast cavity. Philos. Trans. R. Soc. Lond. B Biol. Sci. 377, 20210254 (2022).

13. Kaufmann, P., Black, S. & Huppertz, B. Endovascular trophoblast invasion: implications for the pathogenesis of intrauterine growth retardation and preeclampsia. Biol. Reprod. 69, 1–7 (2003).

14. Carter, A. M. et al. Comparative placentation and animal models: patterns of trophoblast invasion -- a workshop report. Placenta 27 Suppl A, S30–3 (2006).

15. Suckow, M. A., Stevens, K. A. & Wilson, R. P. The Laboratory Rabbit, Guinea Pig, Hamster, and Other Rodents. (Academic Press, 2012).

16. Kapoor, A. & Matthews, S. G. Short periods of prenatal stress affect growth, behaviour and hypothalamo-pituitary-adrenal axis activity in male guinea pig offspring. J. Physiol. 566, 967–977 (2005).

17. Deanesly, R. Implantation and early pregnancy in ovariectomized guinea-pigs. J. Reprod. Fertil. 1, 242–248 (1960).

18. Nikas, G., Ao, A., Winston, R. M. & Handyside, A. H. Compaction and surface polarity in the human embryo in vitro. Biol. Reprod. 55, 32–37 (1996).

19. Meistermann, D. et al. Integrated pseudotime analysis of human pre-implantation embryo single-cell transcriptomes reveals the dynamics of lineage specification. Cell Stem Cell 28, 1625–1640.e6 (2021).

20. Niakan, K. K. & Eggan, K. Analysis of human embryos from zygote to blastocyst reveals distinct gene expression patterns relative to the mouse. Dev. Biol. 375, 54–64 (2013).

21. Gerri, C. et al. Initiation of a conserved trophectoderm program in human, cow and mouse embryos. Nature 587, 443–447 (2020).

22. Wicklow, E. et al. HIPPO pathway members restrict SOX2 to the inner cell mass where it promotes ICM fates in the mouse blastocyst. PLoS Genet. 10, e1004618 (2014).

23. Frum, T., Murphy, T. M. & Ralston, A. HIPPO signaling resolves embryonic cell fate conflicts during establishment of pluripotency in vivo. Elife 7, (2018).

24. Petropoulos, S. et al. Single-Cell RNA-Seq Reveals Lineage and X Chromosome Dynamics in Human Preimplantation Embryos. Cell 165, 1012–1026 (2016).

25. Boroviak, T. et al. Single cell transcriptome analysis of human, marmoset and mouse embryos reveals common and divergent features of preimplantation development. Development 145, (2018).

26. Stirparo, G. G. et al. Integrated analysis of single-cell embryo data yields a unified transcriptome signature for the human pre-implantation epiblast. Development 145, (2018).

27. Posfai, E. et al. Position- and Hippo signaling-dependent plasticity during lineage segregation in the early mouse embryo. Elife 6, (2017).

28. Tsang, M., Friesel, R., Kudoh, T. & Dawid, I. B. Identification of Sef, a novel modulator of FGF signalling. Nat. Cell Biol. 4, 165–169 (2002).

29. Fürthauer, M., Lin, W., Ang, S.-L., Thisse, B. & Thisse, C. Sef is a feedback-induced antagonist of Ras/MAPK-mediated FGF signalling. Nat. Cell Biol. 4, 170–174 (2002).

30. Zhu, P. et al. Single-cell DNA methylome sequencing of human preimplantation embryos. Nat. Genet. 50, 12–19 (2018).

31. Radley, A., Corujo-Simon, E., Nichols, J., Smith, A. & Dunn, S.-J. Entropy sorting of single-cell RNA sequencing data reveals the inner cell mass in the human pre-implantation embryo. Stem Cell Reports 18, 47–63 (2023).

32. Bernardo, A. S. et al. Mammalian embryo comparison identifies novel pluripotency genes associated with the naïve or primed state. Biol. Open 7, (2018).

33. Nowotschin, S. et al. The emergent landscape of the mouse gut endoderm at single-cell resolution. Nature 569, 361–367 (2019).

34. Mohammed, H. et al. Single-Cell Landscape of Transcriptional Heterogeneity and Cell Fate Decisions during Mouse Early Gastrulation. Cell Rep. 20, 1215–1228 (2017).

35. Wehrens, R. & Buydens, L. M. C. Self- and Super-organizing Maps inR: ThekohonenPackage. J. Stat. Softw. 21, (2007).

36. Wehrens, R. & Kruisselbrink, J. Flexible self-organizing maps in kohonen 3.0. J. Stat. Softw. 87, (2018).

37. Kanehisa, M. & Goto, S. KEGG: kyoto encyclopedia of genes and genomes. Nucleic Acids Res. 28, 27–30 (2000).

38. Schultz, R. M. The molecular foundations of the maternal to zygotic transition in the preimplantation embryo. Hum. Reprod. Update 8, 323–331 (2002).

39. Ikeda, S., Sugimoto, M. & Kume, S. Importance of methionine metabolism in morula-to-blastocyst transition in bovine preimplantation embryos. J. Reprod. Dev. 58, 91–97 (2012).

40. Sun, H. et al. Methionine adenosyltransferase 2A regulates mouse zygotic genome activation and morula to blastocyst transition†. Biol. Reprod. 100, 601–617 (2019).

41. Li, J. et al. Metabolic control of histone acetylation for precise and timely regulation of minor ZGA in early mammalian embryos. Cell Discov 8, 96 (2022).

42. Ramos-Ibeas, P. et al. Pluripotency and X chromosome dynamics revealed in pig pre-gastrulating embryos by single cell analysis. Nat. Commun. 10, 500 (2019).

43. Wamaitha, S. E. et al. IGF1-mediated human embryonic stem cell self-renewal recapitulates the embryonic niche. Nat. Commun. 11, 764 (2020).

44. Liu, D. et al. Primary specification of blastocyst trophectoderm by scRNA-seq: New insights into embryo implantation. Sci Adv 8, eabj3725 (2022).

45. Niwa, H. et al. Interaction between Oct3/4 and Cdx2 determines trophectoderm differentiation. Cell 123, 917–929 (2005).

46. Bai, Q. et al. Dissecting the first transcriptional divergence during human embryonic development. Stem Cell Rev Rep 8, 150–162 (2012).

47. Hubert, M. A., Sherritt, S. L., Bachurski, C. J. & Handwerger, S. Involvement of Transcription Factor NR2F2 in Human Trophoblast Differentiation. PLoS One 5, e9417 (2010).

48. Stuart, T. et al. Comprehensive Integration of Single-Cell Data. Cell 177, 1888–1902.e21 (2019).

49. Yanagida, A. et al. Naive stem cell blastocyst model captures human embryo lineage segregation. Cell Stem Cell 28, 1016–1022.e4 (2021).

50. Deng, Q., Ramsköld, D., Reinius, B. & Sandberg, R. Single-cell RNA-seq reveals dynamic, random monoallelic gene expression in mammalian cells. Science 343, 193–196 (2014).

51. Blakeley, P. et al. Defining the three cell lineages of the human blastocyst by single-cell RNA-seq. Development 142, 3151–3165 (2015).

52. Falco, G. et al. Zscan4: a novel gene expressed exclusively in late 2-cell embryos and embryonic stem cells. Dev. Biol. 307, 539–550 (2007).

53. De Iaco, A. et al. DUX-family transcription factors regulate zygotic genome activation in placental mammals. Nat. Genet. 49, 941–945 (2017).

54. Guo, Y., et al. Obox4 promotes zygotic genome activation upon loss of Dux. bioRxiv 2022.07.04.498763 (2024) doi:10.1101/2022.07.04.498763.

55. Leidenroth, A. & Hewitt, J. E. A family history of DUX4: phylogenetic analysis of DUXA, B, C and Duxbl reveals the ancestral DUX gene. BMC Evol. Biol. 10, 364 (2010).

56. Mirzadeh Azad, F., et al. regulates one-carbon metabolism and histone methylation in ground-state pluripotency. Sci Adv 9, eadg7997 (2023).

57. Guo, G. et al. Human naive epiblast cells possess unrestricted lineage potential. Cell Stem Cell 28, 1040–1056.e6 (2021).

58. Cao, Z. et al. Transcription factor AP-2γ induces early Cdx2 expression and represses HIPPO signaling to specify the trophectoderm lineage. Development 142, 1606–1615 (2015).

59. Choi, I., Carey, T. S., Wilson, C. A. & Knott, J. G. Transcription factor AP-2γ is a core regulator of tight junction biogenesis and cavity formation during mouse early embryogenesis. Development 139, 4623–4632 (2012).

60. Kastan, N. et al. Small-molecule inhibition of Lats kinases may promote Yap-dependent proliferation in postmitotic mammalian tissues. Nat. Commun. 12, 1–12 (2021).

61. Nishioka, N. et al. The Hippo signaling pathway components Lats and Yap pattern Tead4 activity to distinguish mouse trophectoderm from inner cell mass. Dev. Cell 16, 398–410 (2009).

62. Chazaud, C., Yamanaka, Y., Pawson, T. & Rossant, J. Early lineage segregation between epiblast and primitive endoderm in mouse blastocysts through the Grb2-MAPK pathway. Dev. Cell 10, 615– 624 (2006).

63. Nichols, J., Silva, J., Roode, M. & Smith, A. Suppression of Erk signalling promotes ground state pluripotency in the mouse embryo. Development 136, 3215–3222 (2009).

64. Yamanaka, Y., Lanner, F. & Rossant, J. FGF signal-dependent segregation of primitive endoderm and epiblast in the mouse blastocyst. Development 137, 715–724 (2010).

65. Roode, M. et al. Human hypoblast formation is not dependent on FGF signalling. Dev. Biol. 361, 358–363 (2012).

66. Piliszek, A., Madeja, Z. E. & Plusa, B. Suppression of ERK signalling abolishes primitive endoderm formation but does not promote pluripotency in rabbit embryo. Development 144, 3719–3730 (2017).

67. Canizo, J. R. et al. A dose-dependent response to MEK inhibition determines hypoblast fate in bovine embryos. BMC Dev. Biol. 19, 13 (2019).

68. Ohhata, T. & Wutz, A. Reactivation of the inactive X chromosome in development and reprogramming. Cell. Mol. Life Sci. 70, 2443–2461 (2013).

69. Sahakyan, A. et al. Human Naive Pluripotent Stem Cells Model X Chromosome Dampening and X Inactivation. Cell Stem Cell 20, 87–101 (2017).

70. Maclary, E. et al. PRC2 represses transcribed genes on the imprinted inactive X chromosome in mice. Genome Biol. 18, 82 (2017).

71. Vallot, C. et al. XACT Noncoding RNA Competes with XIST in the Control of X Chromosome Activity during Human Early Development. Cell Stem Cell 20, 102–111 (2017).

72. Blandau, R. J. Observations on implantation of the guinea pig ovum. Anat. Rec. 103, 19–47 (1949).

73. Sansom, G. S. & Hill, J. P. Observations on the structure and mode of implantation of the blastocyst of Cavia. Trans. Zool. Soc. Lond. 21, 295–354 (1931).

74. Govindasamy, N., Duethorn, B., Oezgueldez, H. O., Kim, Y. S. & Bedzhov, I. Test-tube embryos - mouse and human development in vitro to blastocyst stage and beyond. Int. J. Dev. Biol. 63, 203– 215 (2019).

75. Lüscher, B., Mitchell, P. J., Williams, T. & Tjian, R. Regulation of transcription factor AP-2 by the morphogen retinoic acid and by second messengers. Genes Dev. 3, 1507–1517 (1989).

76. Plath, K. et al. Role of histone H3 lysine 27 methylation in X inactivation. Science 300, 131–135 (2003).

77. Borensztein, M. et al. Xist-dependent imprinted X inactivation and the early developmental consequences of its failure. Nat. Struct. Mol. Biol. 24, 226–233 (2017).

78. Dossin, F. et al. SPEN integrates transcriptional and epigenetic control of X-inactivation. Nature 578, 455–460 (2020).

79. Okamoto, I. et al. Eutherian mammals use diverse strategies to initiate X-chromosome inactivation during development. Nature 472, 370–374 (2011).

80. Zhou, F. et al. Reconstituting the transcriptome and DNA methylome landscapes of human implantation. Nature 572, 660–664 (2019).

81. van den Berg, I. M., Galjaard, R. J., Laven, J. S. E. & van Doorninck, J. H. XCI in preimplantation mouse and human embryos: first there is remodelling…. Hum. Genet. 130, 203–215 (2011).

82. Okamoto, I. et al. The X chromosome dosage compensation program during the development of cynomolgus monkeys. Science 374, eabd8887 (2021).

83. Ton, M.-L. N. et al. An atlas of rabbit development as a model for single-cell comparative genomics. Nat. Cell Biol. 25, 1061–1072 (2023).

84. Canizo, J., Biondic, S., Lenghan, K. V. & Petropoulos, S. Guinea pig preimplantation embryos: Generation, collection, and immunofluorescence. Methods Mol. Biol. (2023) doi:10.1007/7651_2023_488.

85. Vandal, K., Biondic, S., Canizo, J. & Petropoulos, S. Manual dissociation of mammalian preimplantation embryos for single-cell genomics. Methods Mol. Biol. (2023) doi:10.1007/7651_2023_494.

86. Zhao, C. et al. Single-cell multi-omics of human preimplantation embryos shows susceptibility to glucocorticoids. Genome Res. 32, 1627–1641 (2022).

87. Ramos-Ibeas, P. et al. In vitro culture of ovine embryos up to early gastrulating stages. Development 149, (2022).

88. Kagawa, H. et al. Human blastoids model blastocyst development and implantation. Nature 601, 600–605 (2022).

89. Gafni, O. et al. Derivation of novel human ground state naive pluripotent stem cells. Nature 504, 282–286 (2013).

90. Picelli, S. et al. Full-length RNA-seq from single cells using Smart-seq2. Nat. Protoc. 9, 171–181 (2014).

91. Canizo, J., Vandal, K., Biondic, S. & Petropoulos, S. Whole-Mount RNA, Single-Molecule RNA (smRNA), and DNA Fluorescence In Situ Hybridization (FISH) in Mammalian Embryos. Methods Mol. Biol. (2023) doi:10.1007/7651_2023_490.

92. Depreux, F. F., Czech, L. & Whitlon, D. S. Sex Genotyping of Archival Fixed and Immunolabeled Guinea Pig Cochleas. Sci. Rep. 8, 1–7 (2018).

93. Howe, K. L. et al. Ensembl 2021. Nucleic Acids Res. 49, D884–D891 (2021).

94. Kim, D., Paggi, J. M., Park, C., Bennett, C. & Salzberg, S. L. Graph-based genome alignment and genotyping with HISAT2 and HISAT-genotype. Nat. Biotechnol. 37, 907–915 (2019).

95. Liao, Y., Smyth, G. K. & Shi, W. featureCounts: an efficient general purpose program for assigning sequence reads to genomic features. Bioinformatics 30, 923–930 (2014).

96. Pertea, M. et al. StringTie enables improved reconstruction of a transcriptome from RNA-seq reads. Nat. Biotechnol. 33, 290–295 (2015).

97. Pertea, G. & Pertea, M. GFF Utilities: GffRead and GffCompare. F1000Res. 9, (2020).

98. Camacho, C. et al. BLAST+: architecture and applications. BMC Bioinformatics 10, 421 (2009).

99. Wolozin, B. & Ivanov, P. Stress granules and neurodegeneration. Nat. Rev. Neurosci. 20, 649–666 (2019).

100. Hao, Y. et al. Integrated analysis of multimodal single-cell data. Cell 184, 3573–3587.e29 (2021).

101. McCarthy, D. J., Campbell, K. R., Lun, A. T. L. & Wills, Q. F. Scater: pre-processing, quality control, normalization and visualization of single-cell RNA-seq data in R. Bioinformatics 33, 1179– 1186 (2017).

102. Zappia, L. & Oshlack, A. Clustering trees: a visualization for evaluating clusterings at multiple resolutions. Gigascience 7, giy083 (2018).

103. Yu, G., Wang, L.-G., Han, Y. & He, Q.-Y. clusterProfiler: an R package for comparing biological themes among gene clusters. OMICS 16, 284–287 (2012).

104. Tenenbaum, D., Volkening, J. & Maintainer, B. P. KEGGREST. Bioconductor https://bioconductor.org/packages/release/bioc/html/KEGGREST.html (2023).

105. Qiu, X. et al. Reversed graph embedding resolves complex single-cell trajectories. Nat. Methods 14, 979–982 (2017).

106. Cao, J. et al. The single-cell transcriptional landscape of mammalian organogenesis. Nature 566, 496–502 (2019).

107. Korotkevich, G., Sukhov, V. & Sergushichev, A. Fast gene set enrichment analysis. bioRxiv 060012 (2019) doi:10.1101/060012.

108. Fornes, O. et al. JASPAR 2020: update of the open-access database of transcription factor binding profiles. Nucleic Acids Res. 48, D87–D92 (2020).

109. Machlab, D. et al. monaLisa: an R/Bioconductor package for identifying regulatory motifs. Bioinformatics 38, 2624–2625 (2022).

110. Bergmann, S. et al. Spatial profiling of early primate gastrulation in utero. Nature 609, 136–143 (2022).

111. Nakamura, T. et al. A developmental coordinate of pluripotency among mice, monkeys and humans. Nature 537, 57–62 (2016).

112. Dobin, A. et al. STAR: ultrafast universal RNA-seq aligner. Bioinformatics 29, 15–21 (2013).

113. Li, B. & Dewey, C. N. RSEM: accurate transcript quantification from RNA-Seq data with or without a reference genome. BMC Bioinformatics 12, 323 (2011).

114. Haghverdi, L., Lun, A. T. L., Morgan, M. D. & Marioni, J. C. Batch effects in single-cell RNA-sequencing data are corrected by matching mutual nearest neighbors. Nat. Biotechnol. 36, 421–427 (2018).

