## Supplementary Table 6 for "The Guinea Pig: A New Model for Human Preimplantation Development"

| Supplementary Table 6: Primary antibodies used in guinea pig embryos | | | | | |
| --- | --- | --- | --- | --- | --- |
| Protein | **Marker** | **Catalogue number** | **Host (type)** | **Brand** | **Dilution** |
| SOX2 | EPI | 14981182 | Rat (monoclonal) | Thermofisher Scientific | 1/50 |
| NANOG | EPI | AF2729 | Goat (polyclonal) | R&D system | 1/200 |
| OCT4 | EPI | AF1759-SP | Goat (polyclonal) | R&D system | 1/250 |
| GATA6 | PE | AF1700-SP | Goat (polyclonal) | R&D System | 1/100 |
| GATA4 | PE | sc-25310 | Mouse (monoclonal) | Santa Cruz | 1/500 |
| SOX17 | PE | AF1924-SP | Goat (polyclonal) | R&D system | 1/200 |
| Active YAP | TE | ab205270 | Rabbit (monoclonal) | Abcam | 1/200 |
| YAP | TE | sc-101199 | Mouse  (monoclonal) | Santa Cruz | 1/200 |
| GATA3 | TE | CM405A | Mouse (monoclonal) | Biocare Medical | 1/250 |
| CDX2 | TE | MU392A-5UC | Mouse | Biogenex | 1/250 |
| ZO-1 | Tight junction | 33-9100 | Mouse (monoclonal) | Life technologies | 1/300 |
| H3K27me3 | Histone methylation | 3184297 | Rabbit (polyclonal) | Active Motif also sells by Thermofisher Scientific | 1/500 |
| NR2F2 | Mural TE | ab211776 | Rabbit | Abcam | 1/100 |
