## Supplementary Table 7 for "The Guinea Pig: A New Model for Human Preimplantation Development"

**Supplementary Table 7: *In vitro* culture of Guinea pig embryos**

| Medium | Days in culture | Number of embryos | Number (%) of arrested embryos | Number (%) of blastocysts | Notes |
| --- | --- | --- | --- | --- | --- |
| GTL  (human medium) | E1,5 to E3,5 | 4 | 4 (100) | 0 (0) | arrested |
| mR1ECM  (rat medium) | E1,5-E2,5 to E5,5 | 8 | 8 (100) | 0 (0) | arrested |
| M2  (mouse medium) | E3,5 to E5,5 | 3 | 3 (100) | 0 (0) | arrested |
| KSOM  (mouse medium) | E3,5 to E5,5 | 4 | 4 (100) | 0 (0) | arrested |
| N2B27  (blastoids medium) | E3,5 to E5,5 | 3 | 3 (100) | 0 (0) | arrested |
| Global medium  (human medium) | E4,25 to E5,5-E6,5 | 5 | 3 (60) | 2 (40) | delayed |
| RDH  (rabbit medium) | E4 to E6,5 | 6 | 1 (17) | 5 (83) | delayed |
| N2B27  (blastoid medium) | E4-E4,25 to E5,5 | 10 | 3 (30) | 7 (70) | blastocyst at E5,5 |
| M2  (mouse medium) | E5,5 to E6 | 18 | 0 (0) | 18 (100) | briefly cultured from MB to LB |
| GTL  (human medium) | E5,5 to E6.5 | 3 | 3 (100) | 0 (0) | arrested |

Notes: Embryos in GTL medium were cultured at 5 % CO_2_ and 5 % O_2_. All other media at 5 % CO_2_ and 21 % O_2_. In all cases temperature at 37 °C.
